# Ensemble encoding of conditioned fear by prefrontal somatostatin interneurons

**DOI:** 10.1101/2021.07.18.452791

**Authors:** Kirstie A. Cummings, Sabina Bayshtok, Paul J. Kenny, Roger L. Clem

## Abstract

Neurons preferentially activated by learning have been ascribed the unique potential to encode memory. However, it remains unclear which genetically-defined cell types are recruited as part of such an ensemble, or what role discrete subpopulations play in behavior. Here we show that fear conditioning activates a heterogeneous neural ensemble in the medial prefrontal cortex (mPFC), comprised to a large degree of GABAergic interneurons immunoreactive for somatostatin (SST-INs). Using an intersectional genetic approach, we demonstrate that fear learning-activated SST-INs exhibit distinct circuit properties, are preferentially reactivated during memory retrieval, and mediate the expression of defensive freezing. We further show that a rewarding experience, morphine treatment, activates an orthogonal SST-IN population that exerts opposing control over fear. These results outline an important role for discrete GABAergic ensembles in fear memory encoding, and point to an unappreciated capacity for functional specialization among SST-INs.

## Introduction

Stimuli encountered during learning preferentially activate specific neurons. Studies that monitor and manipulate these populations suggest that they play an essential role in memory encoding and may retain a cellular trace of learning that, when reactivated, mediates memory expression (Bocchio et al., 2017; Clem and Schiller, 2016; Josselyn and Tonegawa, 2020). Despite the theoretical importance of sparse neural coding, the cellular composition of memory-related ensembles is largely unexplored, and therefore the functional contribution of specific cell types, defined by gene expression, morphology, or input-output connectivity, remains poorly understood.

The formation of associative fear memory is primarily attributed to glutamatergic projection neurons (PNs), which express a variety of cellular mechanisms for experience-dependent plasticity (Herry and Johansen, 2014; Malenka and Bear, 2004; Ressler and Maren, 2019). However, GABAergic INs also respond to memory-related cues and extensively modulate the function of PNs. In particular, INs containing parvalbumin- (PV-INs), vasoactive intestinal peptide (VIP-INs) and somatostatin (SST-INs) play key roles in orchestrating circuit dynamics underlying memory acquisition and expression (Artinian and Lacaille, 2018; Courtin et al., 2014; Cummings and Clem, 2020; Krabbe et al., 2019; Lucas and Clem, 2017; Wolff et al., 2014; Xu et al., 2019). Synaptic inhibition is thought to facilitate these processes in part through rhythmic entrainment of PN firing (Headley and Pare, 2017). However, evidence suggests that both learning and recall also rely on synaptic interactions between INs that promote PN firing through disinhibition, for example when the recruitment of one IN subtype leads to a corresponding suppression of another (Artinian and Lacaille, 2018; Letzkus et al., 2015; Lucas and Clem, 2018). Because they modulate large groups of PNs, such complex outcomes of GABAergic transmission could endow discrete subsets of INs with unique influence over memory networks. Consequently, an important unanswered question is whether learning recruits specific INs that participate selectively in memory encoding.

Recently, we demonstrated that SST-INs in the prelimbic cortex exhibit increased synaptic efficacy as well as cue-evoked activity after auditory fear conditioning, suggesting their involvement in memory formation (Cummings and Clem, 2020). We further showed that SST-INs control defensive freezing through activation of a distributed brain network, an effect likely orchestrated through the relief of PV-IN-mediated inhibition. Here, using genetic tagging approaches, we investigate whether SST-INs function in memory encoding as part of a discrete neuronal ensemble. Consistent with a stable cellular representation, prelimbic neurons activated at the time of fear conditioning were essential for subsequent memory expression. Contributing prominently to this heterogeneous population are distinct subset of SST-INs that are preferentially reactivated at memory retrieval and selectively promote defensive freezing. Conversely, cellular tagging during morphine treatment, a primarily rewarding experience (Jiang et al., 2021; Le Merrer et al., 2009), delineates an orthogonal population of SST-INs that is not only distinct from, but also functionally opposed to fear-encoding SST-INs at the behavioral level.

## Results

### Genetic capture of a memory-related ensemble of prefrontal neurons

Previous work has identified the dorsomedial mPFC, comprised largely of the prelimbic cortex, as an important site for conditioned stimulus (CS) processing in auditory fear conditioning and a critical substrate for fear expression (Bagur et al., 2021; Burgos-Robles et al., 2009; Corcoran and Quirk, 2007; Courtin *et al.*, 2014; Cummings and Clem, 2020; Dejean et al., 2016; DeNardo et al., 2019; Herry and Johansen, 2014; Karalis et al., 2016; Sotres-Bayon and Quirk, 2010; Xu *et al.*, 2019). To identify neurons selectively recruited by learning in the prelimbic mPFC, we employed a viral genetic approach to permanently tag these cells with a fluorescent reporter. This strategy relied on expression of an estrogen receptor-dependent Cre-recombinase (ERCreER) under the control of the enhanced synaptic activity responsive element (E-SARE), which contains regulatory sequences derived from the arc immediate-early gene (Kawashima et al., 2013). Importantly, recombination of target alleles following induction of ERCreER depends on the presence of the estrogen receptor ligand 4-hydroxytamoxifen (4-OHT), which together with rapid degradation of ERCreER restricts neuronal tagging to a period of several hours following activity.

A viral vector containing the E-SARE-ERCreER construct was co-injected into the mPFC together with vectors encoding a Cre-dependent enhanced yellow fluorescent protein (DIO-eYFP) and synapsin-driven mCherry (hSyn-mCherry), the latter of which served purely as a marker of viral infusion (**Fig. 1A**). Following viral incubation, mice were subjected to auditory fear conditioning, which consisted of paired presentations of an auditory tone (2 KHz, 80 dB, 20 s) that co-terminated with an aversive foot shock (0.7 mA, 2 s). As a behavioral control condition, a subset of animals underwent the same procedure, except that aversive foot shocks were omitted (tones only). Following conditioning or tone presentations, subjects received intraperitoneal injections of either 4-OHT or vehicle (Veh) solution and were returned to their home cages for 2 weeks to allow for the expression of recombinant alleles. At this point they were subjected to a test of memory retrieval in a context distinct from the training arena (**Fig. S1**), followed by cFos immunohistochemistry, to examine the degree to which neurons that were activated during CS-US pairing were specifically reactivated during CS exposure.

**Figure 1.**
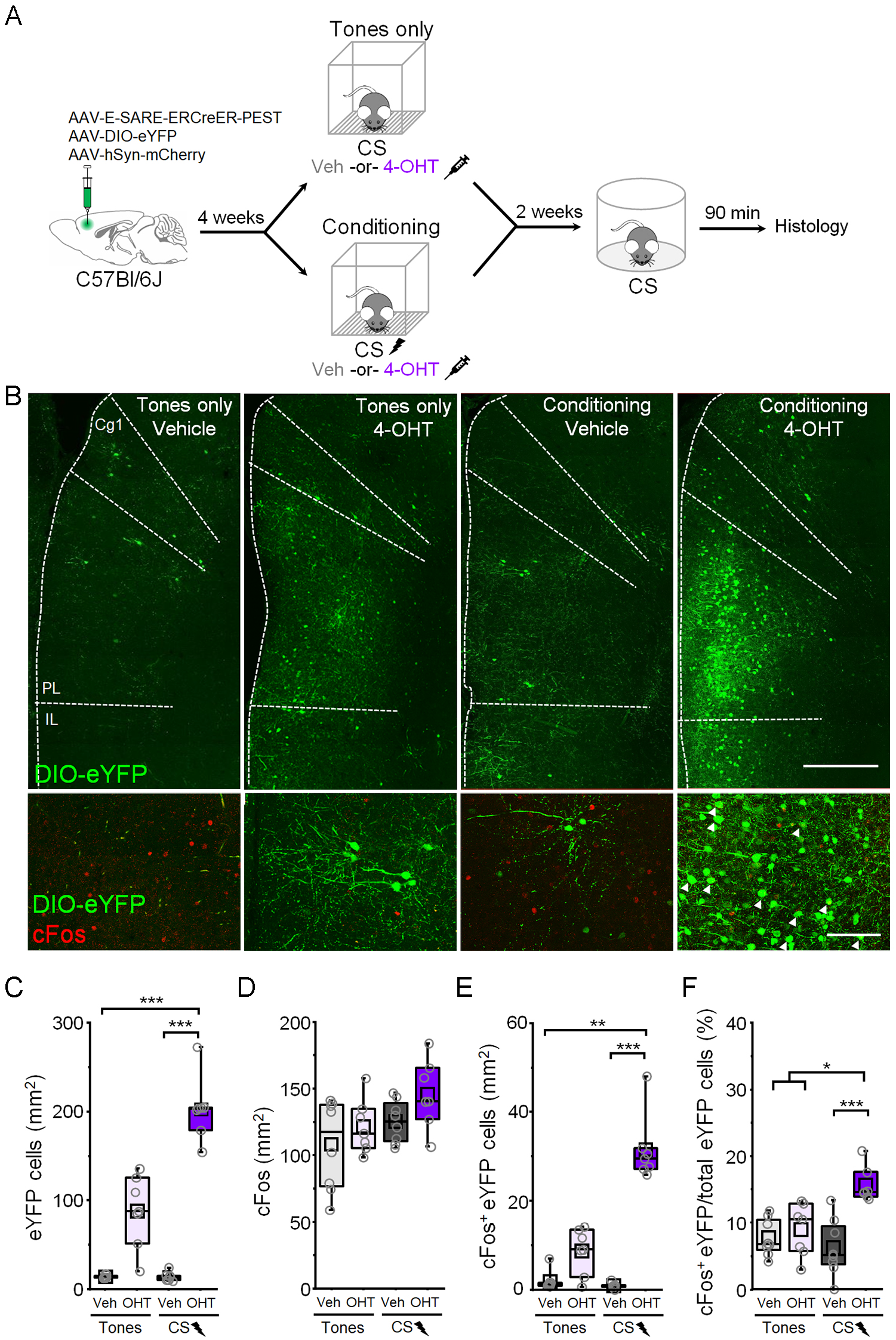
Fear conditioning recruits a prefrontal ensemble that is reactivated upon retrieval. **(A)** Wildtype (C57Bl/6J) mice received bilateral infusions of a cocktail containing vectors encoding E-SARE-ERCreER, Cre-dependent eYFP, and hSyn-mCherry in mPFC. Four weeks later, mice were either subjected to auditory tones (tones only) or CS-US pairing (conditioning) and then immediately injected with vehicle (veh) or 4-hydroxytamoxifen (4-OHT). Two weeks later, in a neutral context, all mice underwent a test of CS-evoked memory retrieval followed by cFos immunohistochemical analysis. **(B)** Top: representative histological images of eYFP tagging. Scale bar = 500 μm. Bottom: Induction of cFos following the retrieval test. White arrowheads denote eYFP tagged neurons immunoreactive for cFos. Scale bar = 100 μm. Cg1 = cingulate area 1. PL = prelimbic cortex. IL = infralimbic cortex. **(C-F)** Comparison between tones only vehicle (n = 8 mice), tones only 4-OHT (n = 7 mice), conditioned vehicle (n = 8 mice), and conditioned 4-OHT (n = 7 mice) groups of **(C)** number of eYFP+ cells: χ^2^ = 23.69 (3), p = 2.89 x 10^−5^, Kruskal-Wallis ANOVA; **(D)** number of cFos+ cells: χ^2^ = 5.75 (3), p = 0.12, Kruskal-Wallis ANOVA; **(E)** number of cFos+/eYFP+ double positive cells: χ^2^ = 20.91 (3), p = 1.09 x 10^−4^, Kruskal-Wallis ANOVA; and **(F)** number of cFos+/eYFP+ double positive cells normalized to the total number of eYFP+ cells: χ^2^ = 16.73 (3), p = 8.05 x 10^−4^, Kruskal-Wallis ANOVA. Experiment was performed in 3 different cohorts and pooled together. * p < 0.05, ** p < 0.01, *** p < 0.001, Dunn’s post hoc test **(C-F)**.

Mice injected with 4-OHT after fear conditioning, but not after tones only exposure, exhibited substantially more eYFP-positive neurons than vehicle-treated mice, which contained only sparse background labeling (**Fig. 1B, C**). Furthermore, while no overall differences in cFos levels were observed following memory retrieval (**Fig. 1D**), mice that received 4-OHT following fear conditioning exhibited a higher density of double-labeled neurons (**Fig. 1E**). More importantly, a higher proportion of eYFP-expressing neurons in these animals were immunoreactive for cFos (**Fig. 1F**), indicating that a greater incidence of double labeling was not attributable to higher levels of eYFP expression. This indicates that neurons activated during fear conditioning are preferentially reactivated during memory retrieval, consistent with their selective involvement in long-term memory.

### Neurons activated during fear learning control memory expression

To test whether learning-activated neurons in the prelimbic cortex mediate memory expression, we next examined whether optogenetic manipulation of these cells influences CS-evoked defensive freezing. To enable optogenetic control of these populations, mice received bilateral infusions of a viral cocktail including E-SARE-ERCreER and hSyn-mCherry vectors, along with a vector encoding either a Cre-dependent Archaerhodopsin (FLEX-Arch3.0-GFP; Arch) or channelrhodopsin construct (DIO-ChR2-eYFP; ChR2), and were implanted with optic fibers directed at prelimbic cortex (**Fig. 2A, B, D, E**). After CS-US pairing, subjects received intraperitoneal injections of either 4-OHT or vehicle and were submitted to a memory retrieval test in which we examined the independent and combined effects of light and CS trials on freezing behavior. Following expression of Arch within learning-activated neurons, photoinhibition (532 nm, 20 s epoch, constant) resulted in a reduction of freezing during CS trials to levels indistinguishable from baseline (**Fig. 2C**). Conversely, after ChR2 expression, photoexcitation (473 nm, 20 s epoch, 20 Hz, 5 ms duration) increased freezing even in the absence of the CS (**Fig. 2F**). No further increase in freezing was observed when photoexcitation was combined with CS presentation, which could potentially reflect a ceiling effect. Examination of difference scores revealed bidirectional effects of optogenetic manipulations during the baseline period (**Fig. 2G**), suggesting that a component of the tagged ensemble might be involved in the expression of contextual fear. The same analysis performed for CS trials revealed a decrease in freezing in 4-OHT-injected Arch-expressing animals compared to all other groups (**Fig. 2H**), suggesting a potential role for the tagged ensemble in cued freezing. Importantly, no light-dependent effects were observed in the vehicle-injected groups, or in animals that received 4-OHT after tones only exposure (**Fig. S2**), indicating that modulation of freezing requires prior conditioning in conjunction with ERCreER activity.

**Figure 2.**
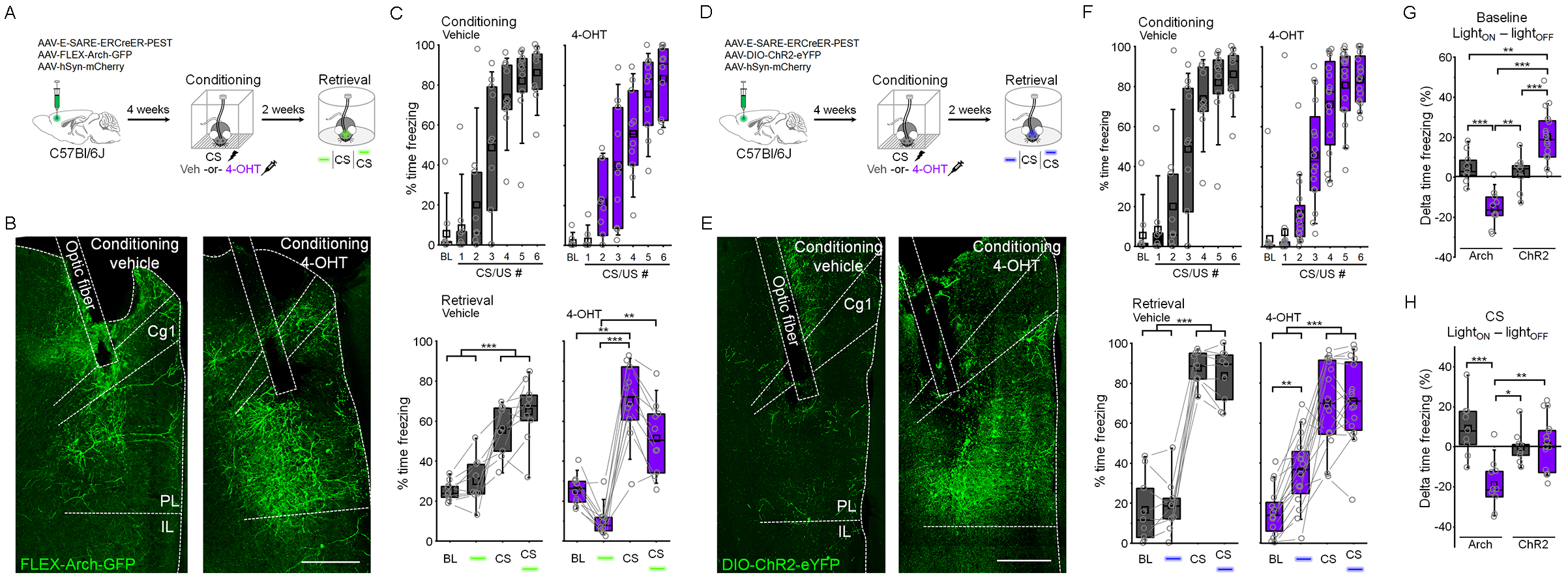
Prefrontal neurons activated by fear learning mediate conditioned freezing. **(A)** For *in vivo* optogenetic silencing of fear learning-related neurons, wildtype mice received bilateral infusions of a cocktail of vectors encoding E-SARE-ERCreER, Cre-dependent Arch, and hSyn-mCherry and were implanted with optic ferrules aimed at PL. Mice were subjected to CS-US pairing and immediately injected with vehicle (veh) or 4-hydroxytamoxifen (4-OHT). Freezing was quantified two weeks later in a neutral context in response to independent and combined light and CS trials. **(B)** Representative histological images of Arch expression and optic fiber placement. Scale = 500 μm. **(C)** Modulation of freezing by light (532 nm solid light, 20 s epochs) and CS trials in vehicle (gray) and 4-OHT (purple) injected mice. Vehicle: F_(1,8)_ = 408.43, p = 3.75 x 10^−8^, 1-way repeated measures ANOVA, n = 9 mice. 4-OHT: χ^2^ = 27.84 (3), p = 3.92 x 10^−6^, Friedman ANOVA, n = 10 mice. Experiments were performed in 3 different cohorts and pooled together. **(D)** For *in vivo* optogenetic activation of fear learning-related neurons, wildtype mice received bilateral infusions of a cocktail of vectors encoding E-SARE-ERCreER, Cre-dependent ChR2, and hSyn-mCherry and were implanted with optic ferrules aimed at PL. Behavior was conducted in a manner identical to **(A)**. **(E)** Representative histological images of ChR2 expression and optic fiber placement. Scale = 500 μm. **(F)** Modulation of freezing by light (473 nm, 5 ms pulses, 20 Hz, 20 s epochs) and CS trials in vehicle (gray) and 4-OHT (purple) injected mice. Vehicle: F_(1,8)_ = 269.4, p = 1.91 x 10^−7^, 1-way repeated measures ANOVA, n = 9 mice. 4-OHT: F_(1,15)_ = 196.72, p = 4.99 x 10^−10^, 1-way repeated measures ANOVA, n = 16 mice. Experiments were performed in 4 different cohorts and pooled together. **(G)** Change in freezing induced by photostimulation during the baseline period in **(C)** and **(F)**. Effect of photostimulation (Light_on_- Light_off_): F_(1,40)_ = 37.16, p = 3.56 x 10^−7^, interaction between opsin and light, 2-way ANOVA. **(H)** Change in freezing induced by photostimulation during CS trials in **(C)** and **(F)**. Effect of photostimulation (Light_on_- Light_off_): F_(1,40)_ = 14.45, p = 4.8 x 10^−4^, interaction between opsin and light, 2-way ANOVA. * p < 0.05, ** p < 0.01, *** p < 0.001, Tukey’s post-hoc test (**c:** vehicle, **F, G**); * p < 0.05, ** p < 0.01, *** p < 0.001, Dunn’s post-hoc test (**c:**4-OHT). Cg1 = cingulate area 1. PL = prelimbic cortex. IL = infralimbic cortex.

### SST-INs are selectively recruited as part of the memory-related ensemble

We recently demonstrated that CS-US pairing increases CS-related activity of prefrontal SST-INs, and that activation of these cells during conditioning is required for subsequent memory expression (Cummings and Clem, 2020). However, it remains unknown whether SST-INs or other GABAergic subtypes participate in memory storage as part of a sparse ensemble. Collectively, PV-INs, VIP-INs and SST-INs comprise >80% of GABAergic neurons in the cortex (Rudy et al., 2011). To determine whether these cell types are recruited into the learning-related ensemble, we performed immunohistochemical staining against markers of these IN subtypes and quantified overlap with an activity-dependent fluorescent tag (**Fig. 3**). In animals that received fear conditioning in combination with 4-OHT, we observed a far greater density of eYFP-positive SST-INs compared to all other conditions (**Fig. 3A-C**). In addition, a higher proportion of tagged neurons in this group expressed SST (**Fig. 3D**), indicating that an increase in labeled SST-INs after conditioning cannot be attributed solely to higher eYFP levels (**Fig. 1**). Importantly, these effects were not observed after tones only exposure, further indicating that labeling of SST-INs is specific to CS-US pairing and not driven by incidental activity. On average, ~20% of the tagged ensemble was comprised of SST-INs (**Fig. 3D**), which in turn represented ~30% of the total SST-IN population (**Fig. 3E**).

**Figure 3.**
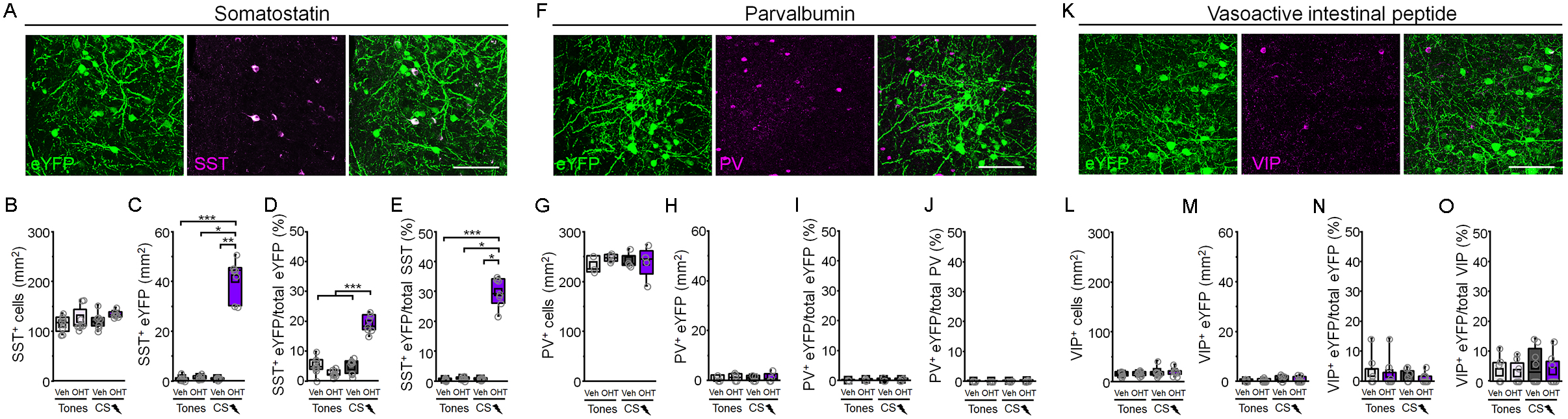
Fear learning-related ensembles are partly comprised of somatostatin-expressing interneurons. **(A)** Immunohistochemical staining against somatostatin (SST) in prelimbic cortex tissue following CS-US pairing and 4-OHT injection. Scale = 50 μm. **(B-E)** Comparison between tones only vehicle (n = 8 mice), tones only 4-OHT (n = 8 mice), conditioning vehicle (n = 8 mice), and conditioning 4-OHT (n = 7 mice) groups of **(B)** SST+ cells, χ^2^ = 6.934 (3), p = 0.074, Kruskal-Wallis ANOVA; **(C)** SST+ eYFP cells, χ^2^ = 16.9 (3), p = 7.4 x 10^−4^, Kruskal-Wallis ANOVA; **(D)** SST+ eYFP cells normalized to the total number of eYFP, F_(1,27)_ = 87.91, p = 5.55 x 10^−10^, interaction between training and treatment, 2-way ANOVA, and **(E)** SST+ eYFP cells normalized to the total number of SST+ cells per group, χ^2^ = 17.04 (3), p = 6.93 x 10^−4^, Kruskal-Wallis ANOVA. **(F)** Immunohistochemical staining against parvalbumin (PV) in prelimbic cortex tissue following CS-US pairing and 4-OHT injection. Scale = 50 μm. **(G-J)** Comparison between tones only vehicle (n = 3), tones only 4-OHT (n = 4), conditioning vehicle (n = 4), and conditioning 4-OHT (n = 4) groups of **(G)** PV+ cells, F_(1,40)_ = 0.73, p = 0.41, 2-way ANOVA; **(H)** PV+ eYFP cells, χ^2^ = 0.485 (3), p = 0.922, Kruskal-Wallis ANOVA; **(I)** PV+ eYFP cells normalized to the total number of eYFP+ cells per group, χ^2^ = 0.882 (3), p = 0.829, Kruskal-Wallis ANOVA; and **(J)** PV+ eYFP cells normalized to the total number of PV+ cells per group, χ^2^ = 1.31 (3), p = 0.727, Kruskal-Wallis ANOVA. **(K)** Immunohistochemical staining against vasoactive intestinal peptide (VIP) in prelimbic cortex tissue following CS-US pairing and 4-OHT injection. Scale = 50 μm. **(L-O)** Comparison between tones only vehicle (n = 8), tones only 4-OHT (n = 8), conditioning vehicle (n = 8), and conditioning 4-OHT (n = 8) groups of **(L)** VIP+ cells, χ^2^ = 1.84 (3), p = 0.606, Kruskal-Wallis ANOVA; **(M)** VIP+ eYFP cells, χ^2^ = 1.56 (3), p = 0.668, Kruskal-Wallis ANOVA; **(N)** VIP+ eYFP cells normalized to the total number of eYFP+ cells per group, χ^2^ = 0.359 (3), p = 0.949, Kruskal-Wallis ANOVA; and **(O)** VIP+ eYFP cells normalized to the total number of VIP+ cells per group, χ^2^ = 1.02 (3), p = 0.796, Kruskal-Wallis ANOVA. Experiments were performed in 2 different cohorts and pooled together. *** p < 0.001, Tukey’s post-hoc test **(D)**. * p < 0.05, ** p < 0.01, *** p < 0.001, Dunn’s post-hoc test **(C,E)**.

In contrast to the above results, learning-activated neurons exhibited negligible immunoreactivity for PV or VIP, and the incidence of recombination among PV- and VIP-INs was not modulated by conditioning or 4-OHT (**Fig. 3F-O**). This could potentially indicate that fear conditioning does not strongly activate these cell types or, alternatively, they may exhibit molecular differences that preclude activation and binding of transcription factors CREB, MEF2 and SRF to E-SARE in response to behavioral experience (Kawashima et al., 2014). While both SST- and VIP-INs respond to aversive foot shocks (Cummings and Clem, 2020; Krabbe *et al.*, 2019; Pi et al., 2013), genetic tagging would require coupling between this type of activity and the above transcriptional responses, which have been linked to memory encoding (Alberini, 2009). Therefore, E-SARE induction potentially demarcates a subset of SST-INs specifically involved in trace formation.

### SST-INs activated during fear learning function like an engram-bearing population

The above results raise the possibility that SST-INs are among the prefrontal neurons activated by learning and subsequently reactivated to mediate memory retrieval (**Fig. 1**), which are important criteria for engram-bearing cells (Tonegawa et al., 2015). Alternatively, activated SST-INs may be relevant only to initial learning, with recall being mediated by other cell types (e.g. glutamatergic PNs). To discriminate between these possibilities, we made use of intersectional activity-dependent tagging to examine the specific role of learning-activated SST-INs. This strategy resembled the one used for non-selective tagging except that in order to be activated by ERCreER, target vectors were also required to undergo flippase (Flp) recombination, which was restricted to SST-INs by the use of SST-FlpO transgenic mice. Prior to training, these animals received bilateral mPFC injections of a viral cocktail including E-SARE-ERCreER and hSyn-mCherry vectors, along with a Cre- and Flp-dependent eYFP construct (Cre_on_/Flp_on_-eYFP) (**Fig. 4A**). Subjects then underwent auditory fear conditioning, or tones only experience (**Fig. S3**), and immediately afterwards received intraperitoneal injections of 4-OHT or vehicle. Following a test of memory retrieval in a context distinct from the training arena, we then utilized cFos immunohistochemistry to examine the degree to which SST-INs that were activated during CS-US pairing were reactivated during CS exposure.

**Figure 4:**
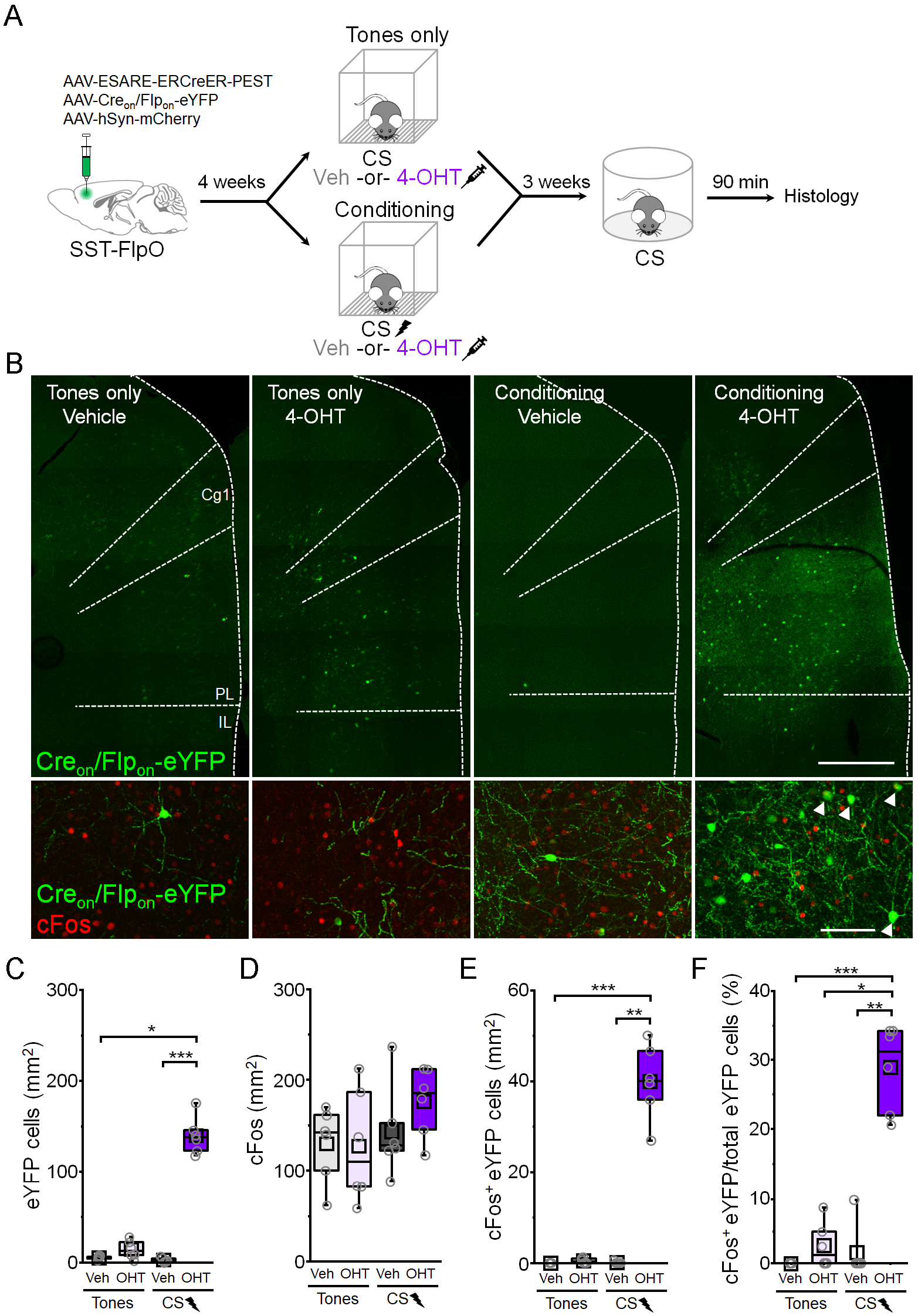
Fear learning-activated somatostatin-interneurons are preferentially reactivated upon memory retrieval. **(A)** SST-FlpO transgenic mice received bilateral infusions into prelimbic cortex of a cocktail containing vectors encoding E-SARE-ERCreER, Cre- and Flp-dependent eYFP, and hSyn-mCherry. Four weeks later, mice received either CS-US pairing (conditioning) or auditory tones (tones only) and were then immediately injected with vehicle (veh) or 4-hydroxytamonifen (4-OHT). After an additional three weeks, in a neutral context, all mice underwent a test of CS-evoked memory retrieval followed by cFos immunohistochemical analysis. **(B)** Top: representative histological images of eYFP tagging. Scale bar = 500 μm. Bottom: Induction of cFos following the retrieval test. White arrowheads denote eYFP neurons immunoreactive for cFos. Scale bar = 100 μm. Cg1 = cingulate area 1. PL = prelimbic cortex. IL = infralimbic cortex. **(C-F)** Comparison between tones only vehicle (n = 6 mice), tones only 4-OHT (n = 6 mice), conditioning vehicle (n = 6 mice), and conditioning 4-OHT (n = 6 mice) groups of **(C)** number of eYFP+ cells: χ^2^ = 17.65 (3), p = 5.2 x 10^−4^, Kruskal-Wallis ANOVA; **(D)** number of cFos+ cells: F_(1,20)_ = 0.82, p = 0.376, 2-way ANOVA; **(E)** number of cFos+/eYFP+ double positive cells: χ^2^ = 17.97 (3), p = 4.46 x 10^−4^, Kruskal-Wallis ANOVA; and **(F)** number of cFos+/eYFP+ double positive cells normalized to the total number of eYFP+ cells in each group: χ^2^ = 17.53 (3), p = 5.49 x 10^−4^, Kruskal-Wallis ANOVA. Experiment was performed in 4 different cohorts and pooled together. * p < 0.05, ** p < 0.01, *** p < 0.001, Dunn’s post-hoc test **(C,E,F)**.

Mice that received 4-OHT after fear conditioning, but not tones only exposure, exhibited a higher density of eYFP-positive cells than vehicle controls (**Fig. 4B, C**). Importantly, we found that a vast majority (>91%) of neurons labeled in this manner are immunoreactive for SST, indicating that our approach captures a relatively pure population of activated SST-INs (**Fig. S4**). Although comparable levels of cFos expression were exhibited across conditions (**Fig. 4D**), conditioning combined with 4-OHT was associated with a higher density of cFos-positive eYFP cells (**Fig. 4E**) and, importantly, a higher proportion of eYFP cells in these animals exhibited cFos immunoreactivity (**Fig. 4F**). This pattern of results suggests that cellular activity during memory retrieval is extraordinarily selective for SST-INs that were active at the time of learning. Indeed, nearly 30% of the tagged SST-IN population was reactivated upon retrieval.

Given these results, we next sought to determine whether SST-INs activated during fear conditioning exert control over memory expression. Prior to training, we injected into the mPFC of SST-FlpO transgenic mice a viral cocktail including E-SARE-ERCreER and hSyn-mCherry vectors, along with a Cre- and Flp-dependent ChR2 construct (Cre_on_/Flp_on_-ChR2-eYFP) (**Fig. 5A-B**). Following auditory fear conditioning, or tones only experience, mice received either 4-OHT or vehicle solution and were submitted to a test of memory retrieval in which we examined the independent and combined effects of light and CS trials. Strikingly, selective photoexcitation (473nm, 20s epochs, 20Hz, 10ms pulses) of a relatively sparse population of tagged SST-INs was on its own sufficient to elicit freezing (**Fig. 5C**), an effect similar to that observed during stimulation of a heterogeneous ensemble (**Fig. 2**). This was not attributable to non-specific motor effects because photoexcitation did not alter locomotor parameters in the open field test (**Fig. S5**). In contrast to these results, mice that received tones only experience or unpaired conditioning prior to SST-IN tagging exhibited no change in freezing upon photoexcitation (**Fig. S6**). These results suggest that fear conditioning activates a subset (~30%) of SST-INs that contribute to the storage and expression of cue associations.

**Figure 5:**
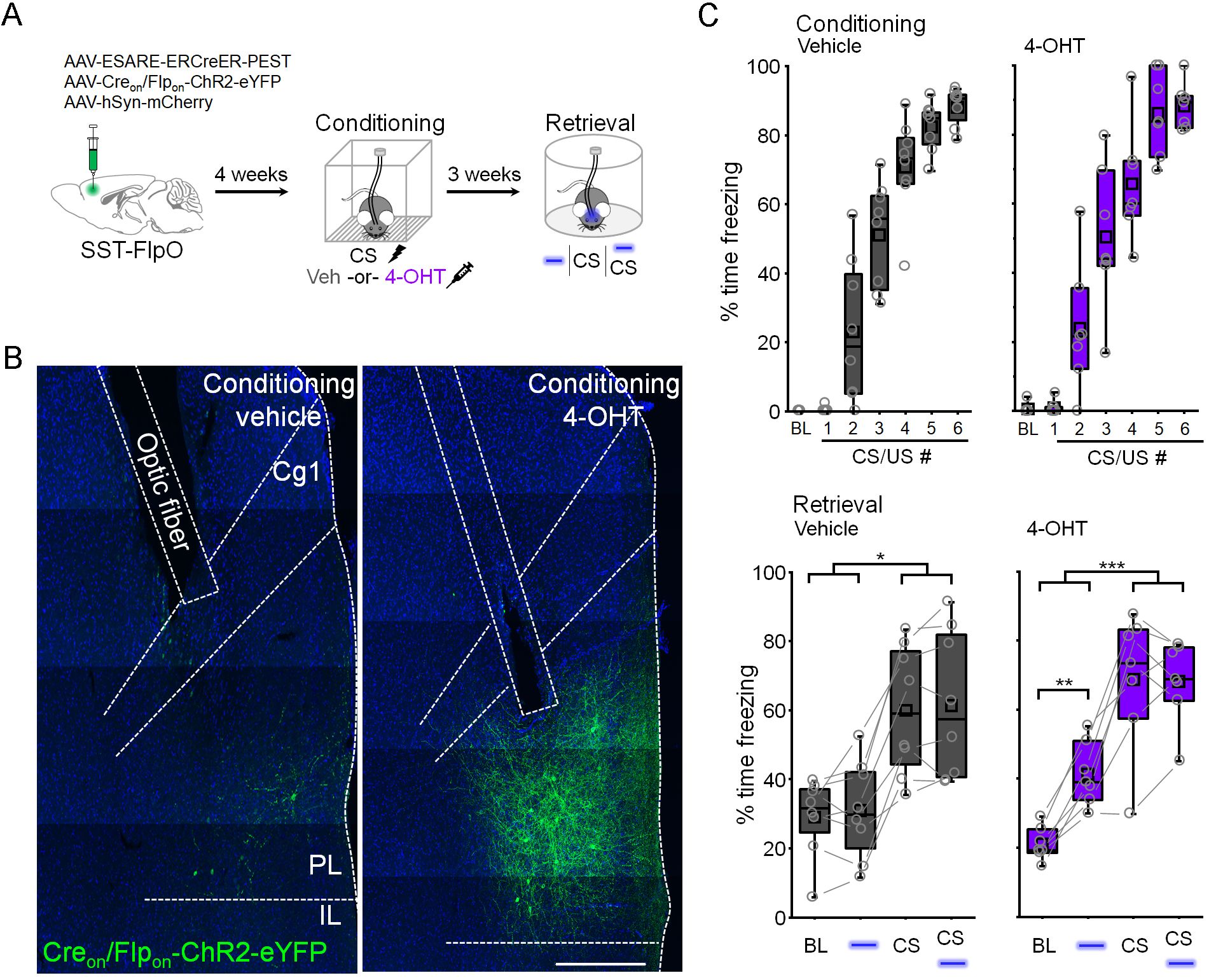
Selective stimulation of learning-activated somatostatin interneurons elicits freezing. **(A)** For *in vivo* optogenetic activation of fear learning-related SST-INs, SST-FlpO transgenic mice received bilateral infusions of a cocktail of vectors encoding E-SARE-ERCreER, Cre- and Flp-dependent ChR2, and hSyn-mCherry and were implanted with optic ferrules aimed at PL. Mice were subjected to CS-US pairing and immediately injected with vehicle (veh) or 4-hydroxytamoxifen (4-OHT). Freezing was quantified three weeks later in a neutral context while testing the independent and combined effect of light and CS trials. **(B)** Representative histological images of ChR2 expression and optic fiber placement. Scale = 500 μm. Cg1 = cingulate area 1. PL = prelimbic cortex. IL = infralimbic cortex. **(C)** Modulation of freezing by light (473 nm, 10 ms pulses, 20 Hz, 20 s epochs) and CS trials in vehicle (gray) and 4-OHT (purple) injected mice. Vehicle: χ^2^ = 19.5 (3), p = 2.15 x 10^−4^, Friedman ANOVA, n = 8 mice. 4-OHT: F_(3,18)_ = 46.7, p = 1.07 x 10^−8^, 1-way repeated measures ANOVA, n = 7 mice. Experiment was performed in 2 different cohorts and pooled together. ** p < 0.01, *** p < 0.001, Tukey’s post-hoc test (**C**: 4-OHT). * p < 0.05, Dunn’s post-hoc test (**C**: vehicle).

### Fear memory-encoding SST-INs exhibit unique input and output synaptic transmission

Memory storage is ultimately thought to be a consequence of synaptic plasticity. Previously, we demonstrated that fear conditioning is associated with an increase in excitatory synaptic transmission onto prelimbic SST-INs, and that SST-IN activation at the time of learning is required both for the expression of this plasticity as well as cue-evoked freezing (Cummings and Clem, 2020). However, it remains unclear whether differences in synaptic transmission can explain the preferential reactivation of learning-activated SST-INs. To examine this possibility, we employed an activity-dependent fluorescent tag in conjunction with constitutive labeling of SST-INs, allowing us to measure the synaptic properties of both tagged and non-tagged SST-INs. First, SST-FlpO x Ai65 transgenic mice received bilateral mPFC infusions of E-SARE-ERCreER and Creon/Flpon-eYFP vectors and then underwent fear conditioning followed by 4-OHT injection (**Fig. 6A**). Electrophysiological recordings were then obtained from learning-activated SST-INs, which were labeled with both eYFP and tdTomato, as well as non-tagged SST-INs, which expressed only tdTomato (**Fig. S7**). SST-INs activated during learning exhibited a higher frequency of spontaneous excitatory postsynaptic currents (EPSCs) compared to non-tagged SST-INs (**Fig. 6B**). However, there were no differences in either frequency or amplitude of inhibitory postsynaptic currents (IPSCs) (**Fig. 6C**). Electrically-evoked EPSCs from tagged SST-INs also exhibited a higher paired-pulse ratio (PPR) compared to those from non-tagged SST-INs, indicating that elevated excitatory transmission onto learning-activated SST-INs is attributable to higher glutamate release probability (**Fig. 6D**). These differences were not observed when comparing tagged and non-tagged SST-INs from animals that received unpaired training (**Fig. S8**), suggesting that they are a specific property of cells engaged in cued associative learning, and given our prior findings, they are likely acquired as part of the memory encoding process (Cummings and Clem, 2020). Interestingly, when we examined action potential firing in response to current injection, we also observed higher excitability of tagged versus non-tagged SST-INs after CS-US pairing specifically in male mice (**Fig. S9**).

**Figure 6:**
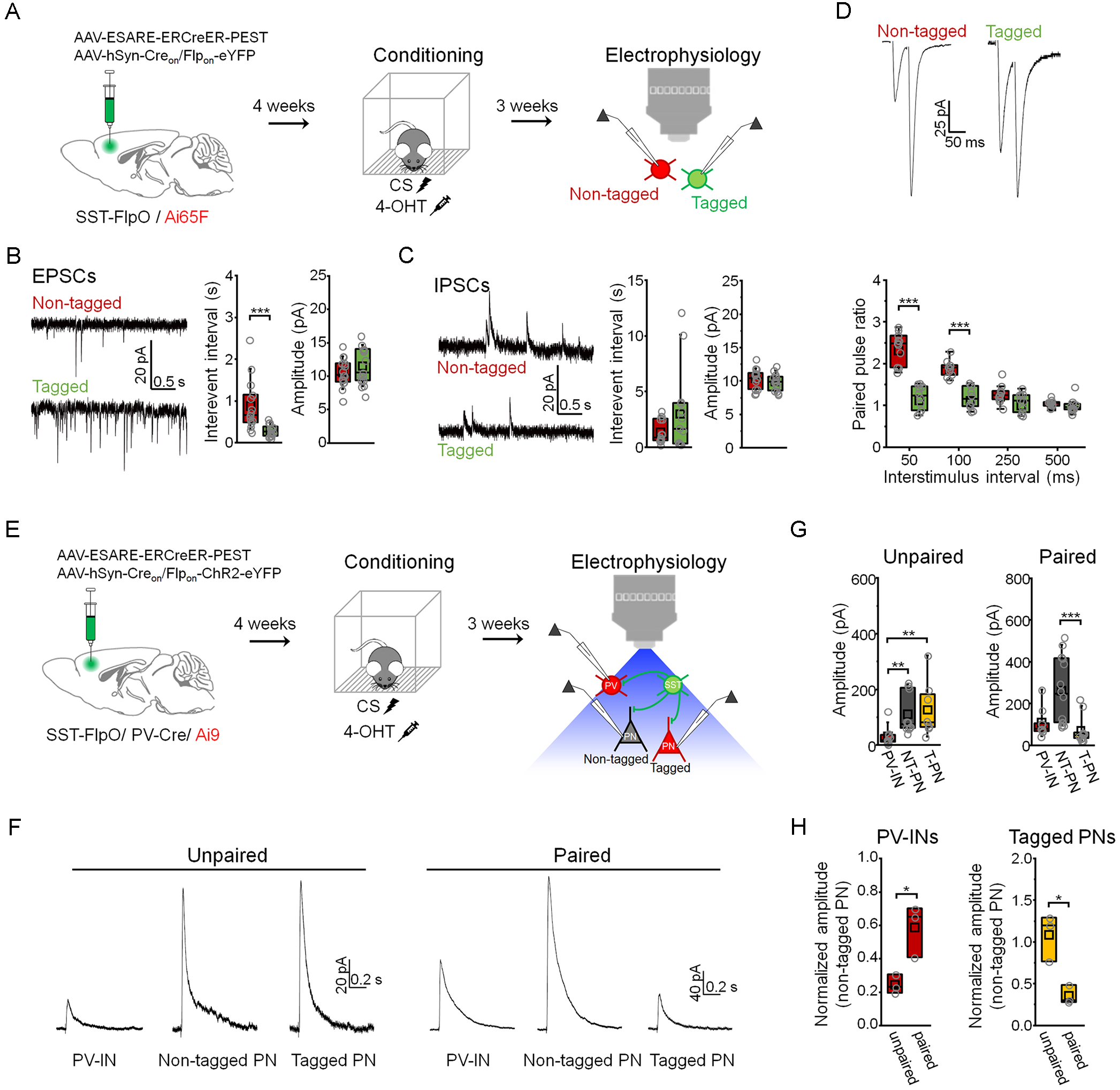
Input and output synaptic connections of learning-activated somatostatin interneurons exhibit distinct properties. **(A)** To independently target and record from both tagged and non-tagged SST-INs, which represent activated and non-activated populations, we infused into the prelimbic cortex of SST-FlpO/ Ai65F double transgenic mice a cocktail of vectors encoding E-SARE-ERCreER, as well as Cre- and Flp-dependent eYFP. Mice were subjected to CS-US pairing and immediately injected with 4-hydroxytamoxifen (4-OHT). Three weeks later, recordings were obtained from eYFP+/ tdTomato+ (tagged) and eYFP−/ tdTomato+ (non-tagged) SST-INs. **(B)** Spontaneous excitatory postsynaptic currents (EPSCs) were collected by interleaved recordings from tagged (n = 15 cells) and non-tagged SST-INs (n = 15 cells) in the same slices (n = 6 slices from 5 mice). Interevent interval: U = 205, p = 1.35 x 10^−4^, Mann-Whitney U-test. Amplitude: t_28_ = −1.03, p = 0.31, two-sided unpaired t-test. **(C)** Spontaneous inhibitory postsynaptic currents (IPSCs) were recorded from tagged (n = 14 cells) and non-tagged SST-INs (n = 14 cells) in the same slices (n = 6 slices from 5 mice). Interevent interval: U = 89, p = 0.696, Mann-Whitney U-test. Amplitude: t_26_ = 0.945, p = 0.353, Mann-Whitney U-test. **(D)** EPSC recordings from tagged (n = 10 cells) and non-tagged SST-INs (n = 11 cells) in the same slices (n = 4 slices in 3 mice) during paired pulse stimulation. Paired pulse ratio: F_(3,7)_ = 16.64, p = 0.0014, interaction between cell type and delay, 2-way repeated measures ANOVA. **(E)** To isolate GABAergic responses at connections from tagged SST-INs onto PV-INs as well as tagged and non-tagged PNs, SST-FlpO/ PV-IRES-Cre/ Ai9 triple transgenic mice received infusions of a cocktail of vectors encoding E-SARE-ERCreER, as well as Cre- and Flp-dependent ChR2. Mice were subjected to either CS-US pairing or unpaired conditioning and injected with 4-OHT. Three weeks later, recordings were obtained from tdTomato+ PV-INs as well as tagged (tdTomato+) and non-tagged (td-Tomato-) PNs. **(F)** Example IPSC traces. **(G)** Amplitude of IPSCs resulting from photoexcitation (460 nm, 1 ms, 0.1 Hz) of tagged SST-INs in PV-INs, non-tagged PNs (NT-PNs), and tagged PNs (T-PNs) following unpaired conditioning (n = 3 slices from 3 mice: PV-INs (n = 10 cells), NT-PNs (n = 10 cells), T-PNs (n = 8 cells)) and CS-US pairing (n = 3 slices from 3 mice: PV-INs (n = 9 cells), NT-PNs (n = 14 cells), T-PNs (n = 12 cells)). Unpaired conditioning: χ^2^ = 12.95 (2), p = 0.0015, Kruskal-Wallis ANOVA. CS-US pairing: χ^2^ = 17.85 (2), p = 1.32 x 10^−4^, Kruskal-Wallis ANOVA. **(H)** IPSC amplitudes of PV-INs and tagged PNs normalized to the median amplitude of non-tagged PNs from the same slices. PV-IN amplitude: t_4_ = 3.56, p = 0.023, two-sided unpaired t-test. Tagged PN amplitude: t_4_ = 4.19, p = 0.013, two-sided unpaired t-test. * p < 0.05, ** p < 0.01, *** p < 0.001 by Mann Whitney U-test (**B**), Tukey’s post-hoc test (**D**), unpaired t-test (**H**) or Dunn’s post-hoc test (**G**).

While increased excitatory input may play a causal role in the reactivation of learning-activated SST-INs, their ability to modulate freezing depends on interaction with other cell types in the mPFC. We previously observed that CS-US pairing, but not unpaired training, alters the balance of transmission from prelimbic SST-INs onto PV-INs versus PNs in a manner that favors PN disinhibition (Cummings and Clem, 2020). We therefore sought to determine whether the microcircuit properties of SST-INs specifically engaged by CS-US pairing differ from those of SST-INs activated under cue non-associative, or unpaired, conditions. Specifically, we examined the relative level of GABAergic inhibition that these populations provide onto neighboring PV-INs as well as PNs that were either tagged or non-tagged as a result of training, in order to reveal how SST-INs interact selectively with learning-activated PNs. To accomplish this, we first generated SST-FlpO x PV-Cre x Ai9 triple transgenic mice, and then bilaterally infused a viral cocktail containing E-SARE-ERCreER and Cre_on_/Flp_on_-ChR2-eYFP vectors into the mPFC of these animals (**Fig. 6E**). Following fear conditioning and 4-OHT injection, this permitted expression of ChR2-eYFP and tdTomato expression in learning-activated SST-INs and PNs, respectively. In addition, constitutive expression of tdTomato was also driven via the PV-Cre allele within PV-INs, which can be readily distinguished from PNs based on electrophysiological properties.

In acute brain slices, we recorded monosynaptic IPSCs in the above cell populations in response to photoexcitation of learning-activated SST-INs (**Fig. 6F**). In unpaired mice, responses of tagged and non-tagged PNs were similar in amplitude, and both cell types exhibited larger responses than PV-INs. In contrast, in paired mice, response amplitudes in tagged PNs were smaller than in non-tagged PNs, while those in non-tagged PNs and PV-INs were equivalent (**Fig. 6G**). To control for potential differences in viral transduction between paired and unpaired animals, IPSCs from PV-INs and tagged PNs were normalized to responses from non-tagged PNs (**Fig. 6H**). This revealed that compared to unpaired training, SST-INs activated during CS-US pairing provide proportionately stronger input onto PV-INs, and generate weaker responses within tagged PNs. These differences in output, which could potentially result from learning-induced plasticity, may facilitate selective disinhibition of memory-related PNs through concerted mono- and disynaptic control.

### Morphine and fear conditioning recruit functionally discrete SST-IN populations

While the above results suggest that memory-encoding SST-INs have distinct circuit properties, it remains possible that freezing elicited by optogenetic stimulation is a non-specific effect of SST-IN transmission, rather than dependent on reactivation of specific cells. In this case, similar effects would be obtained by stimulating other subsets of SST-INs as long as they exceed some threshold population size. Previous work has shown that prelimbic SST-INs exhibit plasticity of membrane and synaptic properties following morphine treatment, an experience highly distinct from fear conditioning (Jiang *et al.*, 2021). We therefore used activity-dependent genetic tagging to investigate whether SST-INs activated by morphine exhibit anatomical overlap with those involved in fear conditioning, and whether reactivation of these cells can drive fear expression after CS-US pairing (**Fig. 7; Fig. S10**). SST-FlpO transgenic mice received viral infusions identical to those employed for fear-related cellular tagging (**Fig. 4**) and were then injected with morphine (10 mg/kg) or saline solution followed 10 hours later by 4-OHT or vehicle (**Fig. 7A**). Three weeks later, all mice underwent CS-US pairing followed by CS presentation and immunohistochemical staining for cFos to determine the degree to which morphine-activated SST-INs were reactivated by fear memory retrieval. As expected, a higher density of eYFP-positive SST-INs was observed following morphine combined with 4-OHT, relative to control conditions (**Fig. 7B-C**). Among the groups, however, a comparable proportion of eYFP-positive SST-INs were immunoreactive for cFos, indicating chance levels of overlap between morphine-related tagging and retrieval-induced activity (**Fig. 7D-F**). This suggests that in contrast to SST-INs activated during fear conditioning (**Fig. 4**), morphine-activated SST-INs were not preferentially reactivated by cued memory retrieval.

**Figure 7:**
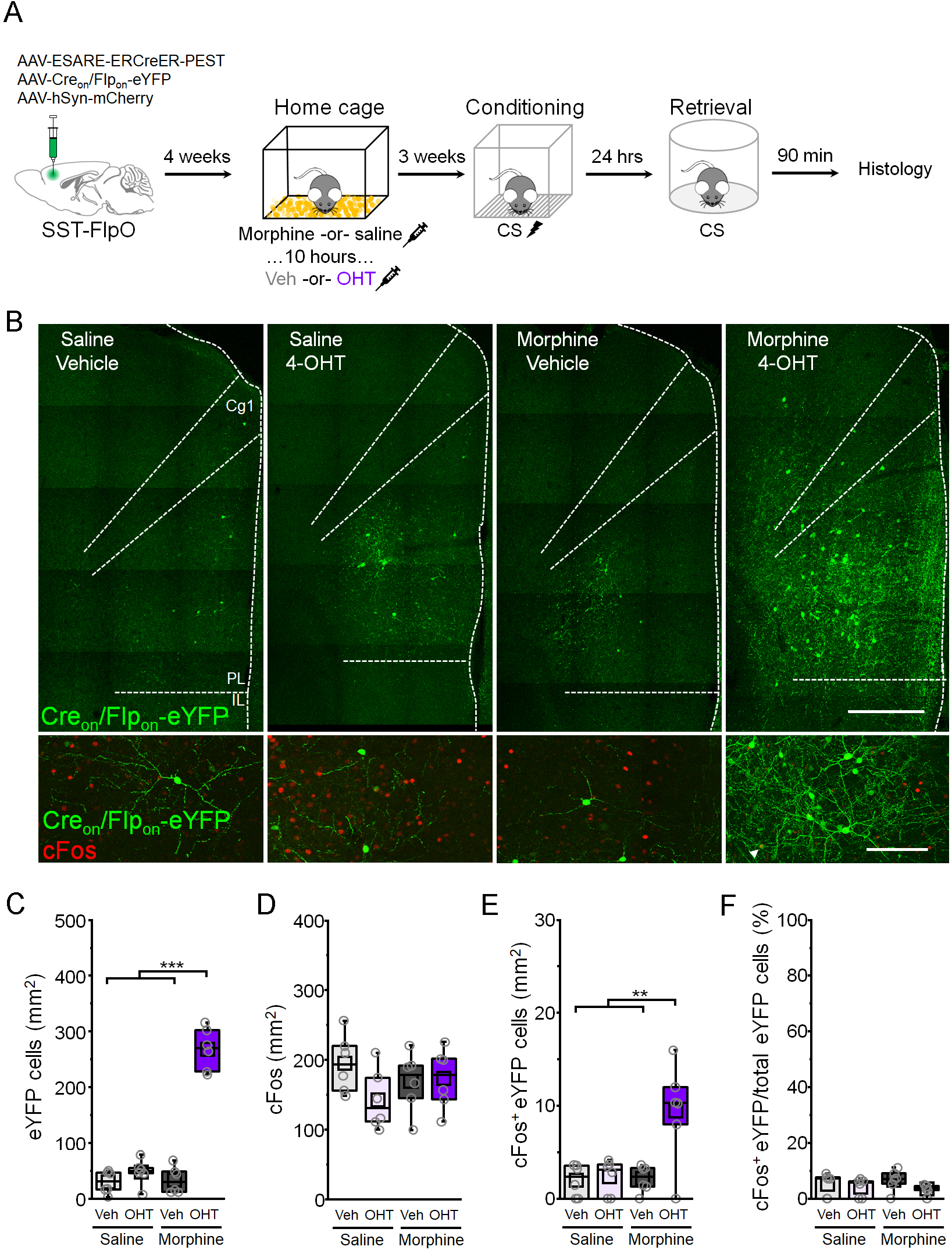
Morphine treatment activates a population of somatostatin-interneurons orthogonal to those involved in fear memory retrieval. **(A)** SST-FlpO transgenic mice received bilateral infusions into prelimbic cortex of a cocktail containing vectors encoding E-SARE-ERCreER, Cre- and Flp-dependent eYFP, and hSyn-mCherry. Four weeks later, in the home cage, they were injected with saline or morphine (10 mg/kg) followed 10 hours later by injections of vehicle (veh) or 4-hydroxytamoxifen (4-OHT). After another 3 weeks, they underwent auditory fear conditioning followed 24 hours later by a test of CS-evoked memory retrieval and cFos immunochemical analysis. (**B**) Top: representative histological images of SST-IN tagging for mice injected with saline and morphine and subsequently injected with either vehicle or 4-OHT. Scale bar = 500 μm. Bottom: Induction of cFos following CS-evoked memory retrieval. White arrowhead denotes cFos+ tagged SST-IN. Scale bar = 100 μm. Cg1 = cingulate area 1. PL = prelimbic cortex. IL = infralimbic cortex. **(C-F)** Comparison between saline vehicle (n = 6 mice), tones only 4-OHT (n = 6 mice), conditioning vehicle (n = 6 mice), and conditioning 4-OHT (n = 6 mice) groups of **(C)** number of eYFP+ cells: F_(1,20)_ = 96.09, p = 4.418 x 10^−9^, interaction between drug and treatment, 2-way ANOVA; **(D)** number of cFos+ puncta: F_(1,20)_ = 2.62, p = 0.121, 2-way ANOVA; **(E)** number of cFos+/eYFP+ double positive cells: F_(1,20)_ = 7.77, p = 0.011, interaction between drug and treatment, 2-way ANOVA; and **(F)** number of cFos+/eYFP+ double positive cells normalized to the total number of eYFP+ cells in each group: χ^2^ = 4.95 (3), p = 0.175, Kruskal-Wallis ANOVA. ** p < 0.01, *** p < 0.001, Tukey’s post-hoc test (**C,E**).

To examine the impact of morphine-activated SST-INs on fear expression, cellular tagging was performed as above, except that E-SARE-ERCreER was used to drive ChR2-eYFP expression to enable subsequent optogenetic manipulation (**Fig. 8A-B**). Three weeks later, photoexcitation of morphine tagged SST-INs (473 nm, 20 s epochs, 20 Hz, 10 ms pulses) had no effect on freezing levels in shock-naïve animals (**Fig. 8C**). On the next day, all subjects underwent CS-US pairing followed by a test of fear memory retrieval in which we examined the independent and combined effects of light and CS trials. In contrast to stimulation of SST-INs that were activated during fear conditioning (**Fig. 5**), photoexcitation of morphine-activated SST-INs had no effect on baseline freezing levels. Interestingly, however, when photoexcitation of morphine-activated SST-INs was combined with CS presentation, freezing levels were reduced compared to CS-only trials (**Fig. 8D**). Importantly, photoexcitation of morphine-activated SST-INs had no effect on open field behavior, implying that reduction of freezing during the test of memory retrieval was not attributable to non-specific locomotor effects (**Fig. S11**). These results indicate that although both morphine and fear conditioning recruit large populations of SST-INs, they exert distinct behavioral effects. Not only do SST-INs activated during fear conditioning uniquely promote fear expression, but they are functionally opposed by morphine-activated SST-INs.

**Figure 8:**
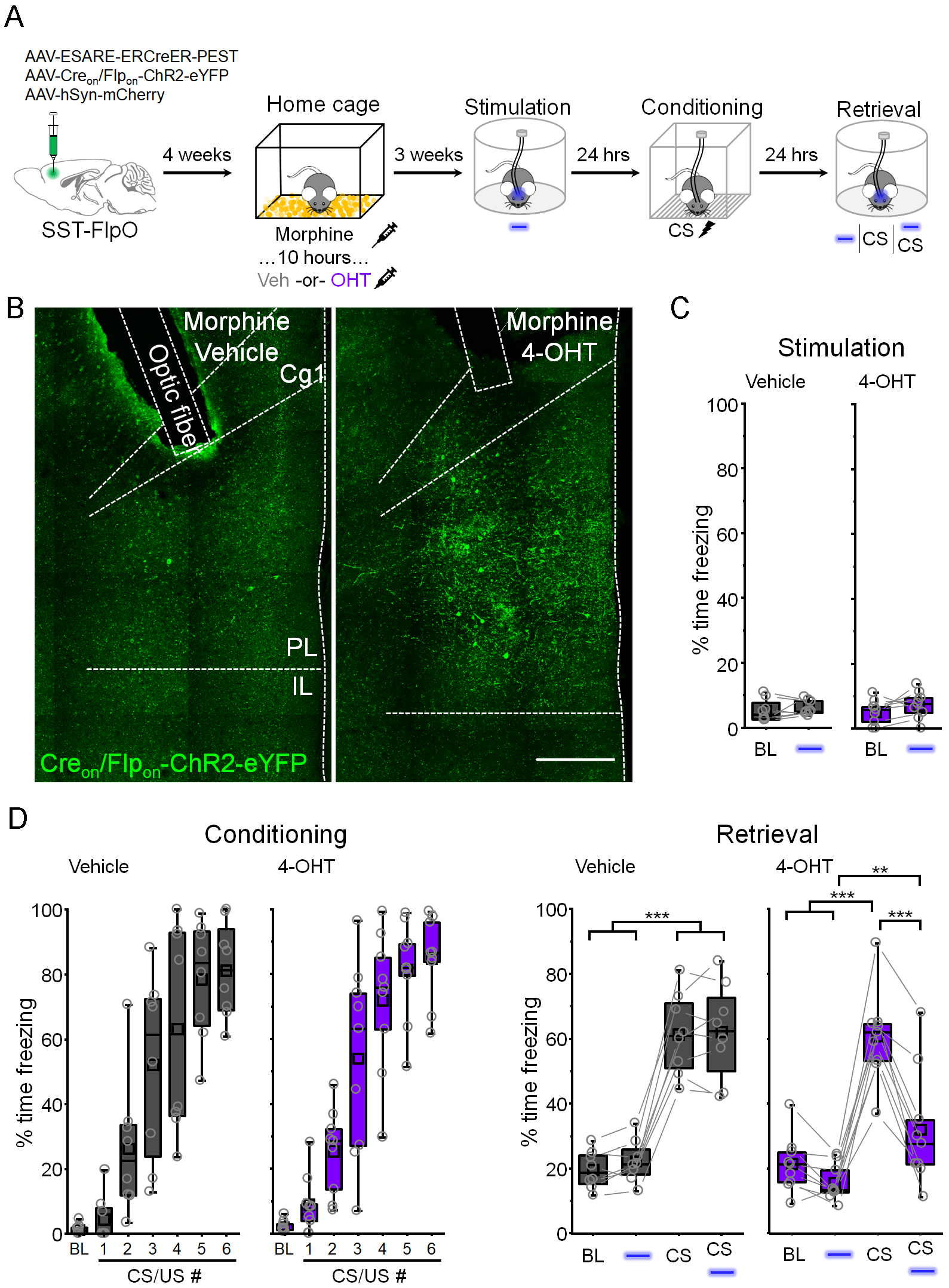
Somatostatin-interneurons activated by morphine oppose conditioned fear expression. **(A)** For *in vivo* optogenetic activation of morphine-related SST-INs, SST-FlpO transgenic mice received bilateral infusions into prelimbic cortex of a cocktail of vectors encoding E-SARE-ERCreER, Cre- and Flp-dependent ChR2, and hSyn-mCherry and were implanted with optic ferrules aimed at PL. Mice received an injection of morphine (10 mg/kg) and 10 hours later, injections of vehicle (veh) or 4-hydroxytamoxifen (4-OHT). Photostimulation-induced freezing (Stimulation) was quantified three weeks later in a neutral context. Mice were then subjected to CS-US pairing (Conditioning) and 24 hours later, a CS-evoked memory retrieval test (Retrieval) in a neutral context, where the independent and combined effects of light and CS trials were quantified. **(B)** Representative histological images of ChR2 expression and optic fiber placement. Scale = 500 μm. Cg1 = cingulate area 1. PL = prelimbic cortex. IL = infralimbic cortex. **(C)** Modulation of freezing by photoexcitation (473 nm, 10 ms pulses, 20 Hz, 20 s epochs) in vehicle (gray) and 4-OHT (purple) mice. Vehicle: t_7_ = −0.788, p = 0.456, paired t-test. 4-OHT: t_8_ = −1.74, p = 0.119, paired t-test. **(D)** Modulation of freezing by photoexcitation (473 nm, 10 ms pulses, 20 Hz, 20 s epochs) and CS presentation during the retrieval test for vehicle (gray) and 4-OHT (purple) mice. Vehicle: F_(3,21)_ = 109.48, p = 5.54 x 10^−13^, 1-way repeated measures ANOVA. 4-OHT: F_(3,24)_ = 46.69, p = 3.59 x 10^−10^, 1-way repeated measures ANOVA. ** p < 0.01, *** p < 0.001, Tukey’s post-hoc test (**D**).

## Discussion

In this study, we demonstrate that a sparse ensemble of neurons in the prelimbic mPFC plays a critical role in fear memory encoding. A major subset of cells activated during learning are SST-INs, whose responsiveness to activity-dependent genetic tagging differentiates them from both PV-INs and VIP-INs. Intersectional genetic capture of learning-activated SST-INs reveals that they are selectively reactivated during memory retrieval and support the expression of defensive freezing, properties that are attributable to differences in transmission within synaptic networks formed by these cells. In particular, memory-related SST-INs receive stronger excitatory input and exhibit biases in GABAergic output that favor disinhibition of PNs recruited by learning. Importantly, fear encoding is a specific property of this cellular subset that is not generalizable to other SST-IN populations. Indeed, SST-INs recruited by a distinct positive experience, morphine exposure, exert opposing control over cued fear expression.

A likely key to the function of memory-encoding SST-INs is that they interact differentially with other cell populations involved in learning. Like SST-INs activated by unpaired conditioning, those activated by CS-US pairing give rise to monosynaptic connections onto PV-INs as well as both activated and non-activated PNs (**Fig. 6**). However, the balance of GABAergic transmission onto these populations differed between the training conditions, reflecting either the recruitment of discrete populations of SST-INs or the expression of plasticity in their output synapses. SST-INs engaged by cued associative learning provide proportionately stronger input onto PV-INs, consistent with previous recordings from our laboratory involving non-selective SST-IN stimulation (Cummings and Clem, 2020). In addition, while SST-INs activated by unpaired training uniformly inhibit PNs, those activated by CS-US pairing provide much weaker inhibition onto activated versus non-activated PNs. The unique network configuration of memory-encoding SST-INs may facilitate PV-IN-mediated disinhibition while simultaneously sparing fear-related PNs from SST-IN-mediated inhibition, enabling recruitment of relevant outputs from the prelimbic cortex while preserving or even augmenting the suppression of irrelevant pathways.

In contrast to SST-INs activated by fear conditioning, those labeled in response to morphine treatment did not elicit freezing either before or after fear conditioning (**Fig. 8**). While this provides important confirmation of the behavioral specificity of memory-related SST-INs, an additional unexpected finding was that stimulation of morphine-related SST-INs opposes the expression of CS-evoked freezing. It is tempting to speculate based on these results that valence is a functional specialization of these discrete populations or otherwise encoded by these cells through experience-dependent plasticity (Tye, 2018). In either case, representation of positive and negative experiences by different subsets of SST-INs raises the possibility that they recruit distinct PNs through disinhibition. In addition, they may compete for behavioral control through mutually opposing inhibition of these output populations (Garcia-Junco-Clemente et al., 2017). Future work should therefore establish whether morphine-activated SST-INs control distinct brain networks or interact directly with cell populations underlying fear memory expression, which may provide unique insights into fear attenuation.

Comparable to prior studies that employed genetic tagging of learning-activated neurons, our results identify a sparse cell population in the prelimbic cortex with predicted attributes of a memory engram (Cai et al., 2016; DeNardo *et al.*, 2019; Han et al., 2009; Kim and Cho, 2017; Lacagnina et al., 2019; Liu et al., 2012; Rashid et al., 2016; Reijmers et al., 2007; Ryan et al., 2015; Tayler et al., 2013). While most reports do not explicitly address the cellular composition of engram populations, or focus exclusively on the role of glutamatergic projections, we demonstrate that fear learning activates a heterogeneous ensemble containing GABAergic cells. Not only do activated SST-INs fulfill conventional criteria for engram-bearing neurons, but their stimulation recapitulates the effect of broader ensemble activation. This suggests that a subset of SST-INs can fully orchestrate network dynamics underlying memory retrieval and likely control the reactivation of glutamatergic PNs. Because SST-INs undergo plasticity during learning, our work underscores the importance of establishing how both excitatory and inhibitory neurons contribute to the formation of a cellular memory trace, as well as how these cell types potentially interact to promote memory expression.

Finally, while a great deal of insight can be gained from parsing the contributions of GABAergic subtypes, functional classification of these cells based on molecular markers is inherently complicated. As one of the most diverse of the broadly defined subtypes, SST-INs have been subclassified based on distinct firing phenotypes, morphologies, and co-expression of secondary markers (Yavorska and Wehr, 2016). For example, while Martinotti cells are estimated to comprise at least half of SST-INs, disinhibition of the thalamorecipient layer of somatosensory cortex is mediated by SST-INs with non-Martinotti morphology (Xu et al., 2013). Within the frontal cortex, functionally relevant subpopulations of SST-INs have been discriminated based on expression of receptors for acetylcholine and oxytocin, as well as signaling components like neuropeptide Y and neuronal nitric oxide synthase (Funk et al., 2017; Jackson et al., 2018; Li et al., 2016; Nakajima et al., 2014; Yamamuro et al., 2020). Interestingly, SST-INs that express oxytocin or nicotinic acetycholine receptors have been implicated in social and anxiety-related behavior (Li *et al.*, 2016; Nakajima *et al.*, 2014; Yamamuro *et al.*, 2020), which are modulated by the processing of aversive stimuli. Nevertheless, our appreciation of how discrete subpopulations encode different aspects of experience is severely limited. Functional mapping of SST-INs, aided by activity-dependent genetic tagging, may help resolve these questions.

In conclusion, we reveal that embedded within a heterogeneous ensemble of neurons underlying associative fear learning are a subset of SST-INs with distinct circuit properties. These cells are preferentially reactivated during memory recall and have a unique capacity to drive fear expression. Our results provide important insight into the cellular specificity of memory encoding by GABAergic microcircuits and outline synaptic mechanisms by which INs may become functionally specialized during memory allocation.

## Author contributions

K.A.C. and R.L.C. initiated the project; R.L.C. and P.J.K. supervised research; K.A.C. and R.L.C. designed experiments; K.A.C. and S.B. performed the research and data analysis; R.L.C. and K.A.C. wrote the manuscript.

## Acknowledgments

We thank Anosha Khawaja for animal care, Haruhiko Bito for sharing the E-SARE-ERCreER plasmid, and members of the Clem laboratory for helpful discussions. These experiments were supported by funds from the National Institute of Mental Health (NIMH) grants R01 MH116145 and R21 MH114170 to R.L.C., in addition to F32 MH115688 and K99 MH122228 to K.A.C.

