## Supplemental materials for "Ensemble encoding of conditioned fear by prefrontal somatostatin interneurons"

### Methods

#### Animals

All procedures were approved by the Institutional Animal Care and Use Committee at the Icahn School of Medicine at Mount Sinai. Male and female mice aged P42-45 at the time of surgery and P70-P98 at the time of behavioral and electrophysiological experiments were used throughout this study. Mice (C57Bl/6J background) were purchased from Jackson Laboratories and housed in groups of 3-5 in a 12 hour light-dark cycle with *ad libitum* access to food and water. Behavioral and electrophysiological experiments were performed during the animals' light cycle. The following genotypes were used: C57Bl/6J (Stock No. 000664), SST-FlpO (Stock No. 028579), PV-IRES-Cre (Stock No. 017320), Ai9 (Stock No. 007909), and Ai65F (Stock No. 032864). Animals were randomly assigned to experimental groups.

#### Vectors

Vectors used in this study that were purchased from Addgene include AAV1-EF1a-DIO-hChR2(H134R)-eYFP-WPRE (Addgene # 20298), AAV1-CBA-FLEX-Arch-GFP (Addgene # 22222), AAV1-EF1a-DIO-eYFP-WPRE (Addgene # 27056), and AAV8-hSyn-mCherry (Addgene # 114472). Vectors purchased from the University of North Carolina Gene Therapy Center Vector Core include rAAVDJ/nEF-$Cre_{on}$/$Flp_{on}$-hChR2(H134R)-eYFP and rAAVDJ/nEF-$Cre_{on}$/$Flp_{on}$-eYFP. The plasmid for the ESARE-ERCreER-PEST vector was a gift from Dr. H. Bito (University of Tokyo). We expanded the plasmid in transformation-competent E. coli, followed by purification using a Qiagen MaxiPrep kit and further extraction in a mixture of phenol-chloroform-isoamyl alcohol (25:24:1) saturated with 10 mM Tris (pH 8.0) and 1 mM EDTA. The purified plasmid was packaged in the AAV8 serotype at the Boston Children's Hospital vector core.

**Stereotaxic surgical procedures**

Surgery was performed as previously described (Cummings and Clem, 2020). Briefly, following induction of anesthesia, mice (P42) were mounted in stereotaxic frames. For activity-dependent tagging in C57Bl/6J mice, vectors were mixed in a ratio of 2:7:1 for ESARE-ERCreER, DIO-eYFP or DIO-ChR2-eYFP or FLEX-Arch-GFP, and hSyn-mCherry, respectively. For intersectional activity-dependent tagging in SST-FlpO mice, vectors were mixed in a ratio of 2:4:0.6 for ESARE-ERCreER, $Cre_{on}$/$Flp_{on}$-ChR2-eYFP or $Cre_{on}$/$Flp_{on}$-eYFP, and hSyn-mCherry, respectively. Vectors were thoroughly mixed just prior to surgery and bilaterally infused into prelimbic cortex (400 nL; AP +1.9, DV −2.0, ML +/− 0.9 at a 10° angle) at a rate of 100 nl/min using motorized injectors (World Precision Instruments). For *in vivo* optogenetics experiments, optic fibers were fabricated as previously described (Cummings and Clem) and bilaterally implanted directly above prelimbic cortex at AP: +1.9, DV: -1.6, ML: +/-0.9 at a 10° angle. Virus was allowed in incubate for 4 weeks prior to behavioral training experiments.

**Fear conditioning and retrieval**

All mice were handled for 3 consecutive days for prior to behavioral testing as described previously (Cummings and Clem, 2020). For optogenetic manipulation experiments, mice were also habituated to the patch cords during handling and were tethered to patch cords for all behavioral tests. Auditory fear conditioning was conducted in sound-attenuating chambers (MedAssociates, St. Albans, VT, USA) and began with a 200 s baseline period followed by presentation of 6 pairings of a neutral auditory tone (CS; 2 kHz, 80 dB, 20 s) with a co-terminating foot shock (US; 0.7 mA, 2 s). CS/US pairings were presented with 80 s interstimulus intervals. For naïve mice, animals were placed in the conditioning arena and exposed to 6 tones (same as CS in conditioning experiments) in the absence of shocks. Unpaired conditioning was performed by placing animals in the conditioning arena and presenting 6 CS tones, then returned to their home cage for 15 min after which they were placed back in the conditioning arena and exposed to 6 US foot shocks.

Two (for tagging in WT mice) or three (intersectional tagging in SST-FlpO mice) weeks after behavioral training and tagging (as described below), mice were subjected to a CS-evoked memory retrieval test in a neutral context. For optogenetic experiments, mice received two laser-only trials, followed by 4 CS presentations alternating with and without simultaneous laser stimulation. CS-only or simultaneous CS/laser presentation order was counterbalanced. The two laser-only epochs, two CS/laser epochs, and two CS-only epochs were each averaged and reported. For trials including optogenetic manipulation of tagged neurons in WT mice, a final transmitted intensity of 7–9 mW for 473 nm (for ChR2; 20 Hz, 5 ms for 20 s epochs) or 532 nm (for Arch; constant light, 20 s) laser-generated (Opto Engine, LLC, Midvale, UT, USA) light was used. For trials including optogenetic activation of tagged SST-INs in SST-FlpO mice, a final transmitted intensity of 7–9 mW for 473 nm (for ChR2; 20 Hz, 10 ms for 20 s epochs) laser light was used. Fear conditioning and retrieval videos for optogenetics experiments were recorded and scored manually by an experimenter blind to the identity of the subject. Videos for untethered mice (used for some cFos experiments and electrophysiology) were analyzed automatically using MedAssociates software. Mice with mistargeted virus and/or misplaced optic fibers were excluded from analyses.

**Morphine administration**

Morphine sulfate was dissolved in sterile saline at 1 mg/mL. Mice received a single 10 mg/kg intraperitoneal injection of morphine or saline in the home cage. Mice receiving an injection of morphine displayed characteristic phenotypes, including morphine-induced tail erection and hyperlocomotion (Lee et al., 1977). Mice were left undisturbed for 10 hours prior to receiving injections of vehicle or 4-hydroxytamoxifen (see below).

**Activity-dependent neural tagging**

Ensemble tagging was performed via a single intraperitoneal (IP) injection of either vehicle or 4-hydroxytamoxifen (4-OHT) immediately following paired fear conditioning, tone exposure in naïve mice, and shock exposure in unpaired mice, or 10 hours after morphine or saline injection. 4-OHT was formulated as previously described (Ye et al., 2016). Briefly, 4-OHT (Sigma H6278) was dissolved in DMSO to a concentration of 40 mg/mL, and further diluted in sterile saline containing 1% Tween-80 to a concentration of 1 mg/mL. Mice were injected at a dose of 10 mg/kg. Vehicle consisted of the same components as the 4-OHT solution, and included 2.5% DMSO and 1% Tween-80 in sterile saline, but lacked 4-OHT. Vehicle was injected at the same volume as the 4-OHT mixture (0.1 cc/10 g body weight). Mice were left undisturbed in their home cages for 24 hours after neural tagging to ensure minimal non-specific recombination.

**Open field test**

SST-FlpO mice with PL SST-INs tagged in response to paired conditioning, unpaired conditioning, naïve tone-exposure, as well as morphine/saline administration were acclimated to the room for 30 minutes before the open field experiment. After tethering implanted ferrules to patch cords, mice were placed in the middle of a 42 cm (length) × 42 cm (width) × 30 cm (height) square open field arena. Movements were detected via 15 infrared beams/detectors on each side. Each test consisted of two alternating and counterbalanced light-on (473 nm, 7–9 mW delivered at 20 Hz, 10 ms pulses) and light-off periods lasting 5 minutes each, for a total of 20 min. Infrared beam breaks and locomotor parameters were quantified using Fusion v5.6 SuperFlex software and reported as averages for each of the two light-on or light-off epochs.

**Immunohistochemistry**

For CS-evoked retrieval cFos experiments in neurons tagged in WT (Figure 1) and SST-FlpO (Figure 4) lines, mice were subjected to 4 CS presentations in a neutral context. For analysis of cFos expression following optogenetic activation of tagged SST-INs in SST-FlpO mice (Figure 4), animals were subjected to 6 photostimulation epochs (473nm, 20 Hz, 10 ms pulses) each lasting 20 s. Following CS presentation or photoexcitation, 90 min later, mice were deeply anesthetized and subjected to transcardial perfusion using phosphate buffered saline (PBS) followed by 4% paraformaldehyde (PFA; pH 7.45). Brains were removed and fixed overnight in 4% PFA before being sectioned at 50 micron thickness on the coronal plane using a Leica VT1000S vibratome. Immunohistochemistry was performed on floating sections. cFos staining was done using a rabbit anti-cFos primary antibody (1:1000; Millipore ABE457). Slices were blocked at room temperature for one hour by using 2% normal goat serum in PBS + 0.3% Tween-20 after which they were incubated in the same solution plus the addition of the primary antibody overnight at 4°C. Slices were then incubated with a secondary goat anti-rabbit antibody conjugated to Alexa 647 (1:500; Jackson Immunoresearch 111-605-045) for two hours at room temperature in PBS containing 2% normal goat serum and 0.3% Tween-20. Staining against somatostatin was performed using a rabbit anti-somatostatin-14 antibody (1:1000; Peninsula Laboratories Cat. # T-4103) in PBS containing 5% BSA and 0.25% Triton X-100 following blocking in the same solution. Staining against parvalbumin (1:1000; Millipore Cat. # MAB1572) and vasoactive intestinal peptide (1:500; Immunostar Cat. # 20077) was performed using a mouse anti-parvalbumin and a rabbit anti-VIP antibody, respectively, in PBS containing 5% normal goat serum and 0.3% Tween-20 after blocking in the same solution. Secondary antibody staining was done using goat anti-rabbit conjugated to Alexa 647 (1:500; Jackson Immunoresearch 111-605-045) for tissue previously stained for SST and VIP. Secondary antibody staining was done using goat anti-mouse conjugated to Alexa 647 (1:500; Jackson Immunoresearch 115-605-003) for tissue previously stained for PV. Slices were mounted with

Prolong Antifade Gold mounting medium with DAPI (Life Technologies, Grand Island, NY, USA) and imaged using an inverted Zeiss 780 confocal microscope running Zen Black software (Carl Zeiss Microscopy, Jena, Germany). eYFP+ tagged neurons, cFos+ puncta, and cells positive for SST, PV, and VIP were quantified manually while blind to experimental condition using the Cell Counter plug-in in ImageJ (NIH, Bethesda, MD, USA).

**Slice electrophysiology**

Following deep anesthetization using inhaled isoflurane, mice were decapitated and brains quickly removed and submerged in ice cold (-2° to -4°C) carbogen-bubbled (95% oxygen, 5% CO2) sucrose cutting solution containing (in mM): 210 sucrose, 26.2 $NaHCO_3$, 11 glucose, 2.5 KCl, 1 $NaH_2PO_4$, 0.5 ascorbate, 4 $MgCl_2$, and 0.5 $CaCl_2$. Coronal sections were cut from mPFC at 300 microns thickness and recovered for 45 min at 35°C in carbogen-bubbled artificial cerebrospinal fluid (ACSF) containing (in mM): 119 NaCl, 26.2 $NaHCO_3$, 11 glucose, 2.5 KCl, 1 $NaH_2PO_4$, 2 $MgCl_2$, and 2 $CaCl_2$. Slices were maintained and used for recordings at room temperature. Whole-cell electrodes (2-5 MΩ) were fabricated from borosilicate glass and filled with internal solution (pH 7.25; 295 mOsmol) containing (in mM): 120 Cs-methanesulfonate, 10 HEPES, 10 Na-phosphocreatine, 8 NaCl, 1 QX-314, 0.5 EGTA, 4 Mg-ATP, and 0.4 Na-GTP for voltage-clamp experiments. An internal solution (pH 7.25; 295-300 mOsmol) containing (in mM): 127.5 K-methanesulfonate, 10 HEPES, 5 KCl, 5 Na-phosphocreatine, 2 $MgCl_2$, 0.6 EGTA, 2 Mg-ATP, and 0.3 Na-GTP was used for current-clamp experiments. Slices were recorded and visualized on an upright microscope equipped with DIC optics and light-emitting diode (LED) - coupled 40X objective. Fluorescence was used to target and record from tagged neurons. In addition, electrophysiological properties were used to confirm cell identities, where interneurons exhibited low capacitance and high membrane resistance, whereas excitatory principal neurons exhibited high capacitance and low input resistance, as well as distinct morphology (prominent apical dendrite and large pyramidal-shaped soma).

Recordings of spontaneous excitatory (EPSCs) and inhibitory (IPSCs) postsynaptic currents were done in standard ACSF by clamping cells at -60 mV and 0 mV, respectively. Each cell was recorded for a duration of 5 minutes in gap-free mode. Electrically-evoked paired-pulse recordings were done by using a bipolar stimulating electrode placed in layer 2 in prelimbic cortex. 10 sweeps were recorded and averaged per cell. For light-evoked monosynaptic inhibitory current recordings, cells were clamped at 0 mV in standard ACSF with the addition of 1 μM TTX (Abcam) and 100 μM 4-aminopyrimidine (Abcam). To stimulate light-evoked transmission, we used a TTL-pulsed microscope objective-coupled LED (460 nm, 20 $mW/mm^2$, 1 ms pulse, Prizmatix). 10 sweeps were recorded and averaged for each cell. Data were acquired at 10 kHz and low-pass filtered at 10 kHz for spontaneous and 3kHz for evoked responses using Multiclamp 700B (Molecular Devices, San Jose, CA, USA) and pClamp 10 software (Molecular Devices).

Excitability recordings were performed in standard ACSF plus the addition of glutamatergic and GABAergic transmission blockers CNQX (10 μM) and picrotoxin (100 μM), respectively. Current injections (500 ms in duration) were performed at -20 pA to +90 pA at 10 pA increments and were delivered at 0.1 Hz. Rheobase was defined as the amount of current needed to fire one action potential.

Cells that exhibited access resistance changes of >20% during the recording, cells that did not meet cell electrophysiological identification criteria (fluorescence, biophysical properties, etc.), or cells that yielded unstable recordings/exhibited unacceptable health (>100pA holding current) were excluded from analyses. All analyses were performed in ClampFit 10 for evoked current and action potential recordings (Molecular Devices) and MiniAnalysis (Synaptosoft, Fort Lee, NJ) for spontaneous current recordings by an experimenter blind to, behavior group, cell type, and sex.

### Quantification and statistical analysis

As preconditions for parametric statistical analysis, we tested the assumptions of normality and homogeneity of variance using the Shapiro-Wilk and Levene's tests, respectively. When these assumptions were not met ($p < 0.05$) non-parametric alternatives were used. Due to the lack of a non-parametric alternative to the 2-way ANOVA, this precluded testing for interactions in some experiments where two factors were present. Following a significant omnibus test, pairwise comparisons among all the groups were used to test for significant differences. All statistical tests, values, and significance levels are specified in the associated figure legends. Box plots depict median (line), mean (open square), quartiles (box), and 10-90% range (whiskers). Individual data points (open circles) are overlaid on box plots. Sample sizes were estimated based on prior electrophysiological and behavioral studies from our laboratory. Statistical analysis and graph construction were done in Graphpad Prism (San Diego, CA) and OriginPRO (OriginLab, Northampton, MA).

**Supplemental figure legends**

**Figure S1, related to Figure 1. Quantification of freezing during training and retrieval for mice in Fig 1. (A)** Freezing during tone presentation and retrieval for mice exposed to tones only. Vehicle retrieval: W = 3, p = 0.583, Wilcoxon signed rank test, n = 8 mice. 4-OHT retrieval: W = 5, p = 1, Wilcoxon signed ranked test, n = 7 mice. **(B)** Freezing during CS-US pairing and CS-evoked retrieval for conditioned mice. Vehicle retrieval: W = 0, p = 0.014, Wilcoxon signed rank test, n = 8 mice. 4-OHT retrieval: $t_6 = -11.43$, $p = 2.68 \times 10^{-5}$, paired t-test, n = 7 mice. * $p < 0.05$, *** $p < 0.001$ by Wilcoxon signed rank (**B**: vehicle) and paired t-test (**B**: 4-OHT). Experiment was performed in 3 different cohorts and pooled together.

**Figure S2 related to Figure 2. No effect of photostimulation of neurons tagged during tones only training. (A)** For *in vivo* optogenetic activation of neurons tagged during tones only training, wildtype mice received bilateral infusions into prelimbic cortex of a cocktail of vectors encoding E-SARE-ERCreER, Cre-dependent ChR2, and hSyn-mCherry and were implantedwith optic ferrules aimed at PL. Mice were presented with 6 auditory tones and immediately injected with vehicle (veh) or 4-hydroxytamoxifen (4-OHT). Freezing was quantified two weeks later in a neutral context while testing the independent and combined effect of light and tone presentation. **(B)** Representative histological images of ChR2 expression and optic fiber placement. Scale = 500 µm. **(C)** Quantification of freezing during photoexcitation (473 nm, 5 ms pulses, 20 Hz, 20 s epochs) and CS presentation in vehicle (gray, n = 7 mice) and 4-OHT (purple, n = 6 mice) mice. Retrieval vehicle: $\chi^2 = 0.214$ (3), p = 0.975, Friedman ANOVA. Retrieval 4-OHT: $\chi^2 = 0.65$ (3), p = 0.885, Friedman ANOVA. Experiment was performed in 2 different cohorts and pooled together.

**Figure S3, related to Figure 4. Quantification of freezing during training and retrieval for SST-FlpO mice. (A)** Freezing during tone presentation and retrieval for mice exposed to tones only. Vehicle retrieval: W = 3, p = 0.361, Wilcoxon signed rank test, n = 6 mice. 4-OHT retrieval: W = 4, p = 0.789, Wilcoxon signed ranked test, n = 6 mice. **(B)** Freezing during CS-US pairing and CS-evoked retrieval for conditioned mice. Vehicle retrieval: W = 0, p = 0.036, Wilcoxon signed rank test, n = 6 mice. 4-OHT retrieval: $t_5 = -9.95$, $p = 1.75 \times 10^{-4}$, paired t-test, n = 6 mice. Experiment was performed in 4 different cohorts and pooled together. *, $p < 0.05$, ***, $p < 0.001$ by Wilcoxon signed rank (**B**: vehicle) and paired t-test (**B**: 4-OHT).

**Figure S4, related to Figure 4. Validation of intersectional tagging in SST-FlpO mice.** For a random subset of 4-OHT-injected conditioned mice (n = 4) from Fig. 4, tissue was stained against somatostatin. The percent of tagged neurons positive for SST was quantified and reported as an average across all mice. Scale = 100 µm. Experiment was performed in 2 different cohorts and pooled together.

**Figure S5, related to Figure 5. Photostimulation of learning-activated SST-INs does not affect locomotion. (A)** SST-FlpO transgenic mice received bilateral infusions into prelimbic cortex of a cocktail containing vectors encoding E-SARE-ERCreER, Cre- and Flp-dependent ChR2, and hSyn-mCherry. Four weeks later, mice were subjected to CS-US pairing and then immediately injected with vehicle (veh) or 4-hydroxytamonifen (4-OHT). After three weeks, locomotion was tested in a 20-minute open field test during light-on and light-off periods (473 nm, 10 ms pulses, 20 Hz). Light on/off epochs were 5 minutes long and were presented in a counterbalanced fashion. **(B)** Example activity plot and quantification of locomotor parameters for 4-OHT mice (n = 6 mice). Distance moved: $t_5 = 0.822$, p = 0.448, paired t-test. Time resting: $t_5 = -0.247$, p = 0.814, paired t-test. Time moving: $t_5 = 0.247$, p = 0.814, paired t-test. **(C)** Example activity plot and quantification of locomotor parameters for vehicle mice (n = 6 mice). Distance moved: $t_5 = 0.724$, p = 0.501, paired t-test. Time resting: $t_5 = 0.795$, p = 0.462, paired t-test. Time moving: $t_5 = -0.793$, $p = 0.463$, paired t-test. Experiments were performed in 2 different cohorts and pooled together.

**Figure S6, related to Figure 5. No effect of photostimulation of SST-INs tagged during tones only or unpaired training in SST-FlpO mice. (A)** For *in vivo* optogenetic activation of SST-INs activated by tones only experience, SST-FlpO transgenic mice received bilateral infusions into prelimbic cortex of a cocktail of vectors encoding E-SARE-ERCreER, Cre- and Flp-dependent ChR2, and hSyn-mCherry and were implanted with optic ferrules aimed at PL. Mice were exposed to 6 auditory tones and immediately injected with vehicle (veh) or 4-hydroxytamoxifen (4-OHT). Freezing was quantified two weeks later in a neutral context while testing the independent and combined effect of light and tone presentation. Scale = 500 µM**. (B)** Modulation of freezing by photoexcitation (473 nm, 5 ms pulses, 20 Hz, 20 s epochs) and tone presentation in vehicle (gray) and 4-OHT (purple) injected mice. Vehicle retrieval: $F_{(3,15)} = 0.712$, $p = 0.559$, 1-way repeated measures ANOVA, n = 6 mice. 4-OHT retrieval: $\chi^2 = 2.61$ (3), $p = 0.455$, Friedman ANOVA, n = 7 mice. Experiment was performed in 2 different cohorts and pooled together. **(C)** For *in vivo* optogenetic activation of SST-INs activated by unpaired conditioning, SST-FlpO transgenic mice underwent surgery as described in (**A**). After 4 weeks, mice underwent unpaired conditioning followed immediately by  injections of vehicle (veh) or 4-hydroxytamoxifen (4-OHT). Freezing was quantified two weeks later in a neutral context while testing the independent and combined effect of light and CS presentation. Scale = 500 µm. **(D)** Quantification of freezing during photoexcitation (473 nm, 5 ms pulses, 20 Hz, 20 s epochs) and CS presentation in vehicle (gray) and 4-OHT (purple) injected mice. Vehicle retrieval: $\chi^2 = 1.99$ (3), $p = 0.575$, Friedman ANOVA, n = 8 mice. 4-OHT retrieval: $\chi^2 = 1.4$ (3), $p = 0.706$, Friedman ANOVA, n = 9 mice. Experiment was performed in 3 different cohorts and pooled together.

**Figure S7, related to Figure 6. Intersectional tagging in SST-FlpO/ Ai65F mice for electrophysiological analysis in Fig. 6.** Representative image illustrating discrimination of tagged (eYFP+) from non-tagged (tdTomato+) SST-INs in the same slice. Scale = 100 µm.

**Figure S8, related to Figure 6. No differences in synaptic input between tagged and non-tagged SST-INs in unpaired mice. (A)** SST-FlpO/ Ai65F double transgenic mice received prelimbic infusions of a cocktail of vectors encoding E-SARE-ERCreER, as well as Cre- and Flp-dependent eYFP. Mice were subjected to unpaired conditioning and immediately injected with 4-hydroxytamoxifen (4-OHT). Three weeks later, recordings were obtained from eYFP+/ tdTomato+ (tagged) and eYFP-/ tdTomato+ (non-tagged) SST-INs. **(B)** Spontaneous excitatory postsynaptic currents (EPSCs) were recorded from tagged (n = 10 cells) and non-tagged SST-INs (n = 10 cells) in the same slices (n = 4 slices from 4 mice). Interevent interval: $t_{18}$ = -0.062, p = 0.95, two-sided unpaired t-test. Amplitude: U = 51, p = 0.97, Mann-Whitney U-test. **(C)** Spontaneous inhibitory postsynaptic currents (IPSCs) were recorded from tagged (n = 10 cells) and non-tagged SST-INs (n = 10 cells) in the same slices (n = 4 slices from 4 mice). Interevent interval: $t_{18}$ = -0.269, p = 0.79, two-sided unpaired t-test. Amplitude: $t_{18}$ = -0.243, p = 0.81, two-sided unpaired t-test. **(D)** EPSC recordings from non-tagged (n = 10 cells) and tagged SST-INs (n = 11 cells) in the same slices (n = 3 slices in 3 mice) during paired pulse stimulation. Paired pulse ratio: $F_{(3,27)}$ = 0.545, p = 0.655, 2-way repeated measures ANOVA.

**Figure S9, related to Figure 6. Comparison of intrinsic excitability between tagged and non-tagged SST-INs of male and female mice after fear conditioning. (A)** SST-FlpO/ Ai65F double transgenic mice received prelimbic infusions of a cocktail of vectors encoding E-SARE-ERCreER, as well as Cre- and Flp-dependent eYFP. Mice were subjected to CS-US pairing and were immediately injected with 4-hydroxytamoxifen (4-OHT). Three weeks later, recordings were obtained from eYFP+/ tdTomato+ (tagged) and eYFP-/ tdTomato+ (non-tagged) SST-INs.

**(B)** Representative spike trains (-20 pA and +10-40 pA current injections), input-output curves, and rheobase quantification for tagged (n = 7 cells) and non-tagged (n = 7 cells) SST-INs in the same slices of male mice (n = 3 slices from 3 mice). Input-output: $F_{(9,36)} = 1.81$, p = 0.1, 2-way repeated measures ANOVA. Rheobase: U = 44, p = 0.009, Mann-Whitney U-test. **(C)** Representative spike trains (-20 pA and +10-40 pA current injections), input-output curves, and rheobase quantification for tagged (n = 14 cells) and non-tagged (n = 16 cells) SST-INs in the same slices of female mice (n = 3 slices from 3 mice). Input-output: $F_{(9,99)} = 0.522$, p = 0.855, 2-way repeated measures ANOVA. Rheobase: U = 126, p = 0.564, Mann-Whitney U-test. **, p < 0.01 by Mann-Whitney U-test (**B**: rheobase).

**Figure S10, related to Figure 7. Quantification of freezing for morphine-treated mice in Fig 7. (A)** Freezing during conditioning and memory retrieval for mice injected with saline. Saline vehicle retrieval: $t_5 = -5.94$, p = 0.002, paired t-test, n = 6 mice. Saline 4-OHT retrieval: $t_5 = -5.94$, p = 0.002, paired t-test, n = 6 mice. **(B)** Freezing during conditioning and memory retrieval for mice injected with morphine. Morphine vehicle retrieval: $t_5 = -9.35$, $p = 2.36 \times 10^{-4}$, paired t-test, n = 6 mice. Morphine 4-OHT retrieval: $t_5 = -5.4$, p = 0.003, paired t-test, n = 6 mice. Experiment was performed in 2 different cohorts and pooled together. **, p < 0.01, ***, p < 0.001 by paired t-test.

**Figure S11, related to Figure 8. Photostimulation of morphine-activated SST-INs does not affect locomotion. (A)** SST-FlpO transgenic mice received bilateral prelimbic infusions of a cocktail containing vectors encoding E-SARE-ERCreER, Cre- and Flp-dependent ChR2, and hSyn-mCherry. Four weeks later, mice were injected with morphine (10 mg/kg) followed 10 hours later by injections of vehicle (veh) or 4-hydroxytamonifen (4-OHT). After three weeks, locomotion was tested in a 20-minute open field test during light-on and light-off periods (473 nm, 10 ms pulses, 20 Hz). Light on/off epochs were 5 minutes long and were presented in a counterbalanced fashion. **(B)** Example activity plot and quantification of locomotor parameters for 4-OHT mice (n = 8 mice). Distance moved: $t_7 = 0.639$, p = 0.543, paired t-test. Time resting: W = 16, p = 0.833, Wilcoxon signed rank test. Time moving: $t_7 = -0.159$, p = 0.878, paired t-test. **(C)** Example activity plot and quantification of locomotor parameters for vehicle mice (n = 8 mice). Distance moved: $t_7 = 1.14$, p = 0.292, paired t-test. Time resting: W = 18, p = 1, Wilcoxon signed rank test. Time moving: W = 18, p = 1, Wilcoxon signed rank test. Experiments were performed in 2 different cohorts and pooled together.

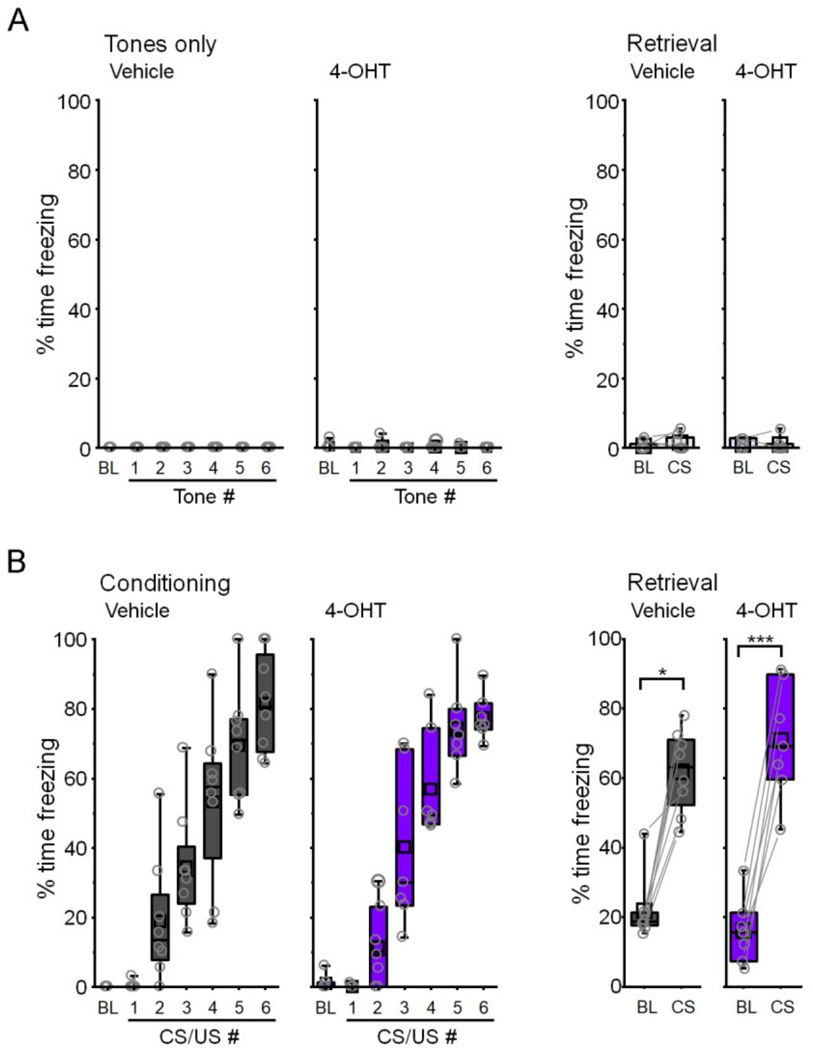


**Figure S1, related to Figure 1. Quantification of freezing during training and retrieval for mice in Fig 1. (A)** Freezing during tone presentation and retrieval for mice exposed to tones only. Vehicle retrieval: W = 3, p = 0.583, Wilcoxon signed rank test, n = 8 mice. 4-OHT retrieval: W = 5, p = 1, Wilcoxon signed ranked test, n = 7 mice. **(B)** Freezing during CS-US pairing and CS-evoked retrieval for conditioned mice. Vehicle retrieval: W = 0, p = 0.014, Wilcoxon signed rank test, n = 8 mice. 4-OHT retrieval: $t_6 = -11.43$, $p = 2.68 \times 10^{-5}$, paired t-test, n = 7 mice. * $p < 0.05$, *** $p < 0.001$ by Wilcoxon signed rank (**B**: vehicle) and paired t-test (**B**: 4-OHT). Experiment was performed in 3 different cohorts and pooled together.

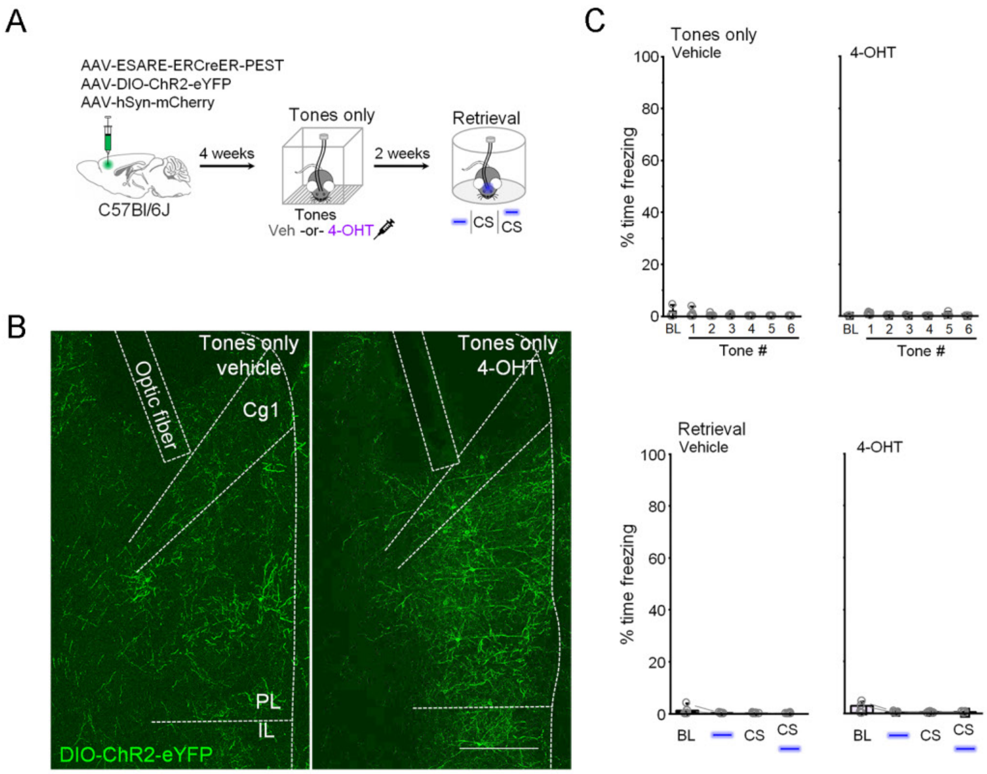


**Figure S2 related to Figure 2. No effect of photostimulation of neurons tagged during tones only training. (A)** For *in vivo* optogenetic activation of neurons tagged during tones only training, wildtype mice received bilateral infusions into prelimbic cortex of a cocktail of vectors encoding E-SARE-ERCreER, Cre-dependent ChR2, and hSyn-mCherry and were implantedwith optic ferrules aimed at PL. Mice were presented with 6 auditory tones and immediately injected with vehicle (veh) or 4-hydroxytamoxifen (4-OHT). Freezing was quantified two weeks later in a neutral context while testing the independent and combined effect of light and tone presentation. **(B)** Representative histological images of ChR2 expression and optic fiber placement. Scale = 500 μm. **(C)** Quantification of freezing during photoexcitation (473 nm, 5 ms pulses, 20 Hz, 20 s epochs) and CS presentation in vehicle (gray, n = 7 mice) and 4-OHT (purple, n = 6 mice) mice. Retrieval vehicle: $\chi^2$ = 0.214 (3), p = 0.975, Friedman ANOVA. Retrieval 4-OHT: $\chi^2$ = 0.65 (3), p = 0.885, Friedman ANOVA. Experiment was performed in 2 different cohorts and pooled together.

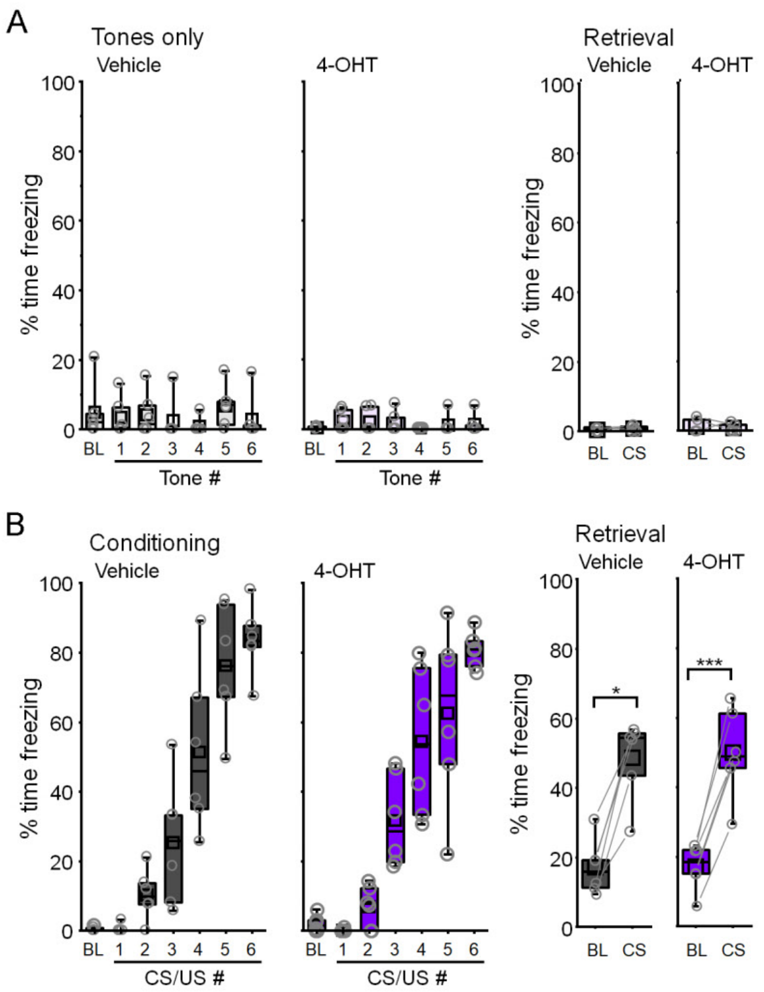


**Figure S3, related to Figure 4. Quantification of freezing during training and retrieval for SST-FlpO mice. (A)** Freezing during tone presentation and retrieval for mice exposed to tones only. Vehicle retrieval: W = 3, $p = 0.361$, Wilcoxon signed rank test, n = 6 mice. 4-OHT retrieval: W = 4, $p = 0.789$, Wilcoxon signed ranked test, n = 6 mice. **(B)** Freezing during CS-US pairing and CS-evoked retrieval for conditioned mice. Vehicle retrieval: W = 0, $p = 0.036$, Wilcoxon signed rank test, n = 6 mice. 4-OHT retrieval: $t_5 = -9.95$, $p = 1.75 \times 10^{-4}$, paired t-test, n = 6 mice. Experiment was performed in 4 different cohorts and pooled together. *, $p < 0.05$, ***, $p < 0.001$ by Wilcoxon signed rank (**B**: vehicle) and paired t-test (**B**: 4-OHT).

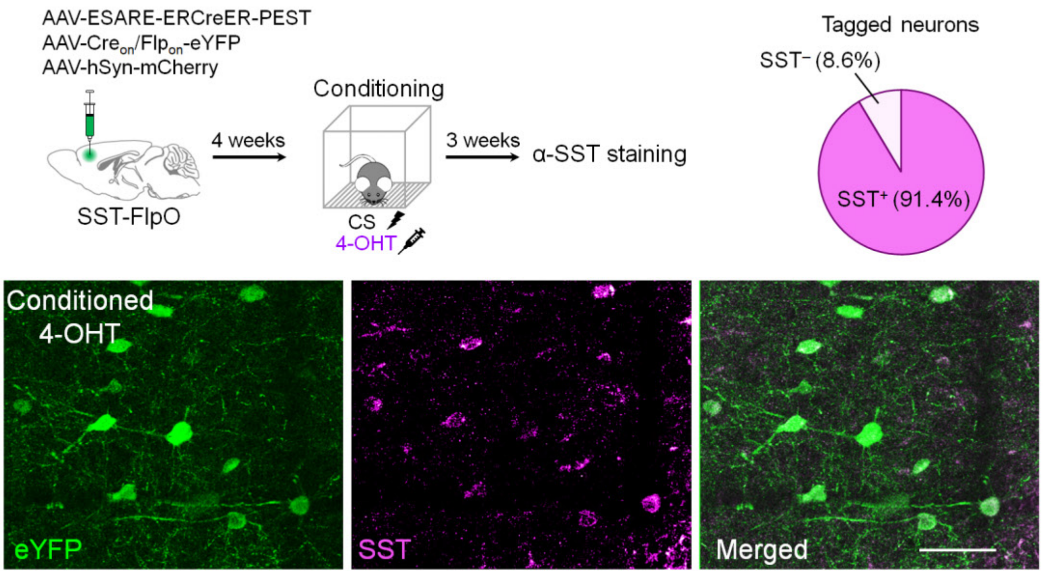


**Figure S4, related to Figure 4. Validation of intersectional tagging in SST-FlpO mice.** For a random subset of 4-OHT-injected conditioned mice (n = 4) from Fig. 4, tissue was stained against somatostatin. The percent of tagged neurons positive for SST was quantified and reported as an average across all mice. Scale = 100 µm. Experiment was performed in 2 different cohorts and pooled together.

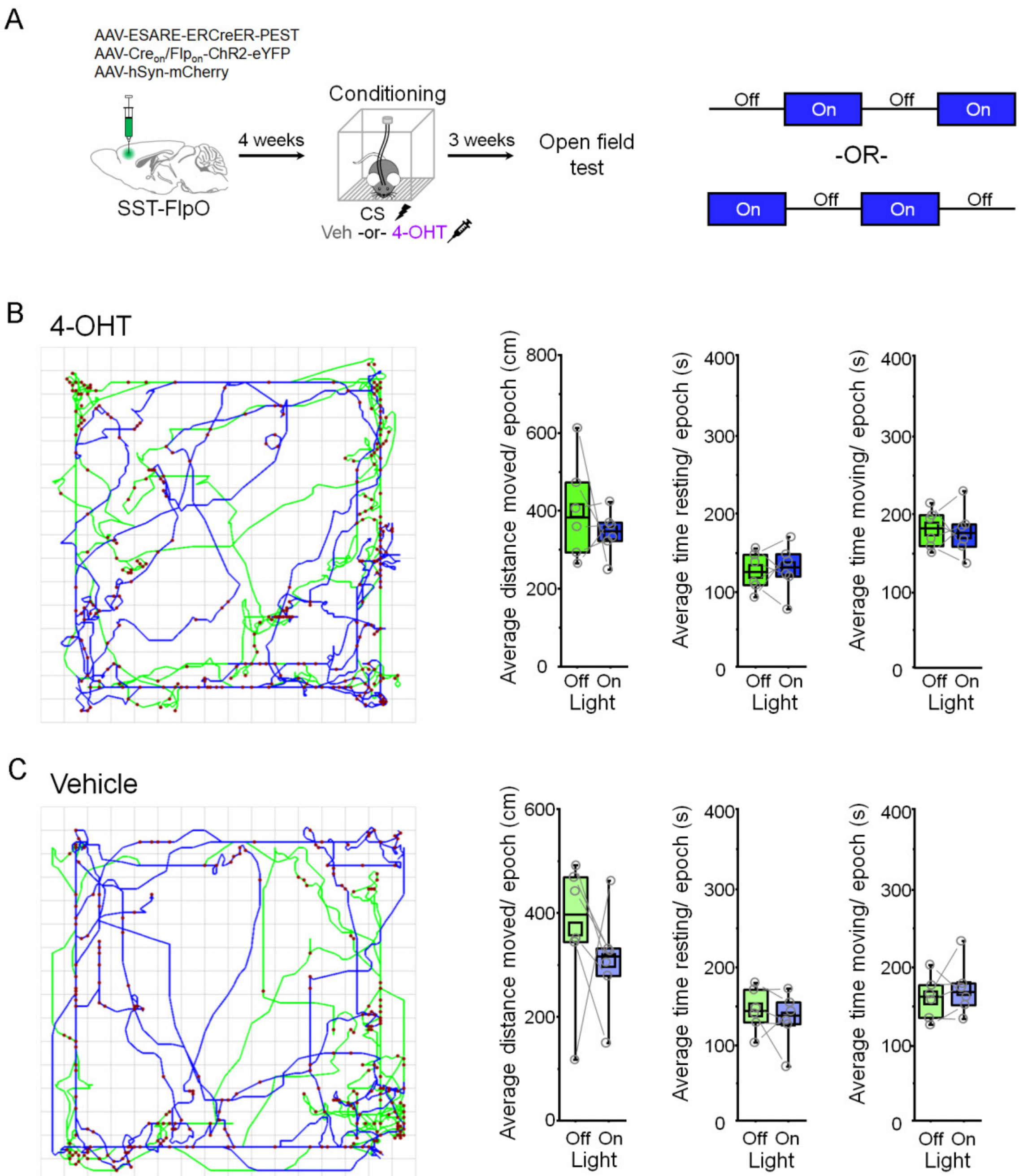


**Figure S5, related to Figure 5. Photostimulation of learning-activated SST-INs does not affect locomotion. (A)** SST-FlpO transgenic mice received bilateral infusions into prelimbic cortex of a cocktail containing vectors encoding E-SARE-ERCreER, Cre- and Flp-dependent ChR2, and hSyn-mCherry. Four weeks later, mice were subjected to CS-US pairing and then immediately injected with vehicle (veh) or 4-hydroxytamonifen (4-OHT). After three weeks, locomotion was tested in a 20-minute open field test during light-on and light-off periods (473 nm, 10 ms pulses, 20 Hz). Light on/off epochs were 5 minutes long and were presented in a counterbalanced fashion. **(B)** Example activity plot and quantification of locomotor parameters for 4-OHT mice (n = 6 mice). Distance moved: $t_5 = 0.822$, $p = 0.448$, paired t-test. Time resting: $t_5 = -0.247$, $p = 0.814$, paired t-test. Time moving: $t_5 = 0.247$, $p = 0.814$, paired t-test. **(C)** Example activity plot and quantification of locomotor parameters for vehicle mice (n = 6 mice). Distance moved: $t_5 = 0.724$, $p = 0.501$, paired t-test. Time resting: $t_5 = 0.795$, $p = 0.462$, paired t-test. Time moving: $t_5 = -0.793$, $p = 0.463$, paired t-test. Experiments were performed in 2 different cohorts and pooled together.

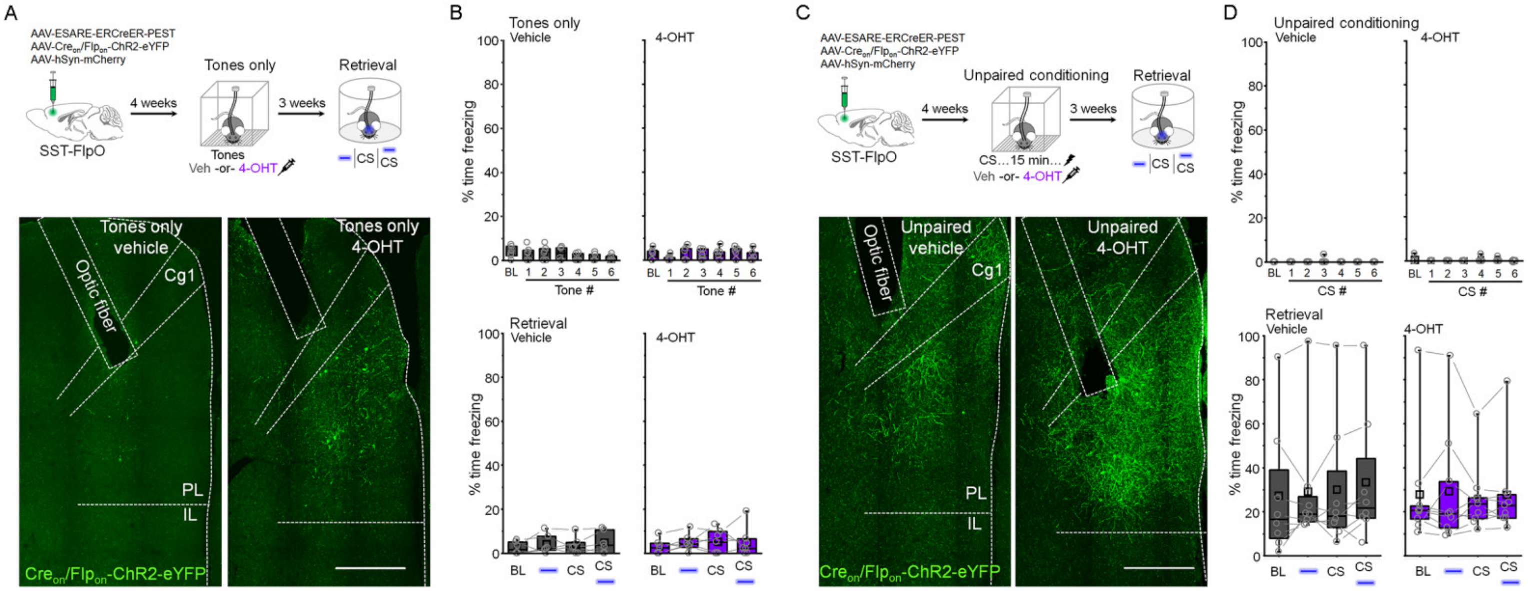

**Figure S6, related to Figure 5. No effect of photostimulation of SST-INs tagged during tones only or unpaired training in SST-FlpO mice. (A)** For *in vivo* optogenetic activation of SST-INs activated by tones only experience, SST-FlpO transgenic mice received bilateral infusions into prelimbic cortex of a cocktail of vectors encoding E-SARE-ERCreER, Cre- and Flp-dependent ChR2, and hSyn-mCherry and were implanted with optic ferrules aimed at PL. Mice were exposed to 6 auditory tones and immediately injected with vehicle (veh) or 4-hydroxytamoxifen (4-OHT). Freezing was quantified two weeks later in a neutral context while testing the independent and combined effect of light and tone presentation. Scale = 500 μM**. (B)** Modulation of freezing by photoexcitation (473 nm, 5 ms pulses, 20 Hz, 20 s epochs) and tone presentation in vehicle (gray) and 4-OHT (purple) injected mice. Vehicle retrieval: $F_{(3,15)} = 0.712$, $p = 0.559$, 1-way repeated measures ANOVA, n = 6 mice. 4-OHT retrieval: $\chi^2 = 2.61$ (3), $p = 0.455$, Friedman ANOVA, n = 7 mice. Experiment was performed in 2 different cohorts and pooled together. **(C)** For *in vivo* optogenetic activation of SST-INs activated by unpaired conditioning, SST-FlpO transgenic mice underwent surgery as described in (**A**). After 4 weeks, mice underwent unpaired conditioning followed immediately by injections of vehicle (veh) or 4-hydroxytamoxifen (4-OHT). Freezing was quantified two weeks later in a neutral context while testing the independent and combined effect of light and CS presentation. Scale = 500 μm. **(D)** Quantification of freezing during photoexcitation (473 nm, 5 ms pulses, 20 Hz, 20 s epochs) and CS presentation in vehicle (gray) and 4-OHT (purple) injected mice. Vehicle retrieval: $\chi^2 = 1.99$ (3), $p = 0.575$, Friedman ANOVA, n = 8 mice. 4-OHT retrieval: $\chi^2 = 1.4$ (3), $p = 0.706$, Friedman ANOVA, n = 9 mice. Experiment was performed in 3 different cohorts and pooled together.

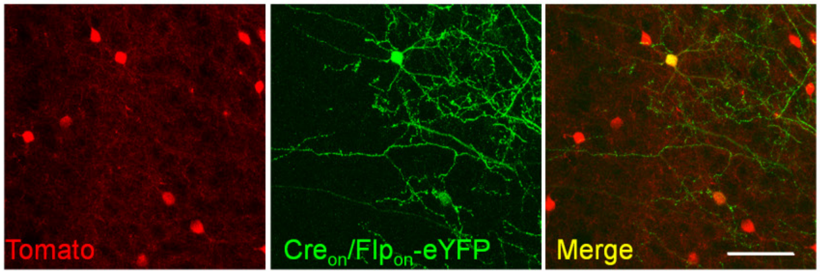


**Figure S7, related to Figure 6. Intersectional tagging in SST-FlpO/ Ai65F mice for electrophysiological analysis in Fig. 6.** Representative image illustrating discrimination of tagged (eYFP+) from non-tagged (tdTomato+) SST-INs in the same slice. Scale = 100 µm.

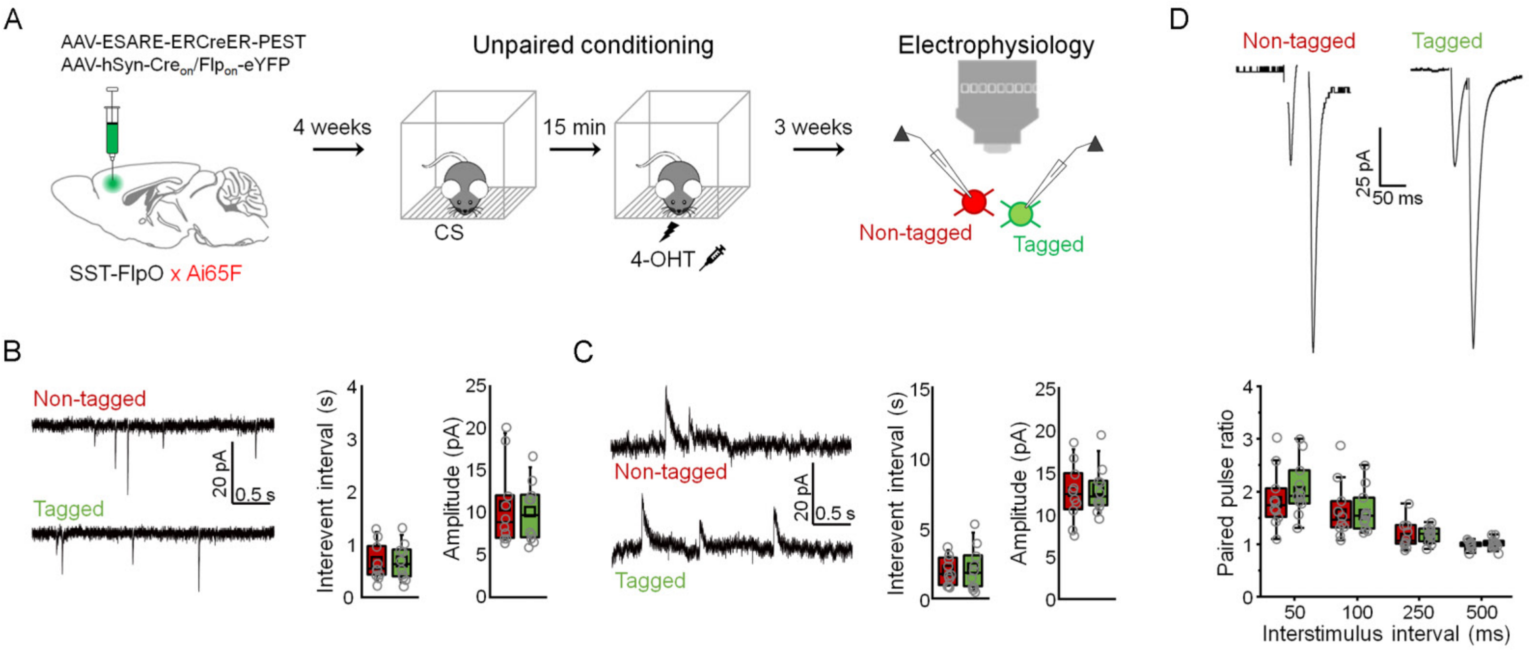


**Figure S8, related to Figure 6. No differences in synaptic input between tagged and non-tagged SST-INs in unpaired mice. (A)** SST-FlpO/ Ai65F double transgenic mice received prelimbic infusions of a cocktail of vectors encoding E-SARE-ERCreER, as well as Cre- and Flp-dependent eYFP. Mice were subjected to unpaired conditioning and immediately injected with 4-hydroxytamoxifen (4-OHT). Three weeks later, recordings were obtained from eYFP+/ tdTomato+ (tagged) and eYFP-/ tdTomato+ (non-tagged) SST-INs. **(B)** Spontaneous excitatory postsynaptic currents (EPSCs) were recorded from tagged (n = 10 cells) and non-tagged SST-INs (n = 10 cells) in the same slices (n = 4 slices from 4 mice). Interevent interval: $t_{18}$ = -0.062, p = 0.95, two-sided unpaired t-test. Amplitude: U = 51, p = 0.97, Mann-Whitney U-test. **(C)** Spontaneous inhibitory postsynaptic currents (IPSCs) were recorded from tagged (n = 10 cells) and non-tagged SST-INs (n = 10 cells) in the same slices (n = 4 slices from 4 mice). Interevent interval: $t_{18}$ = -0.269, p = 0.79, two-sided unpaired t-test. Amplitude: $t_{18}$ = -0.243, p = 0.81, two-sided unpaired t-test. **(D)** EPSC recordings from non-tagged (n = 10 cells) and tagged SST-INs (n = 11 cells) in the same slices (n = 3 slices in 3 mice) during paired pulse stimulation. Paired pulse ratio: $F_{(3,27)}$ = 0.545, p = 0.655, 2-way repeated measures ANOVA.

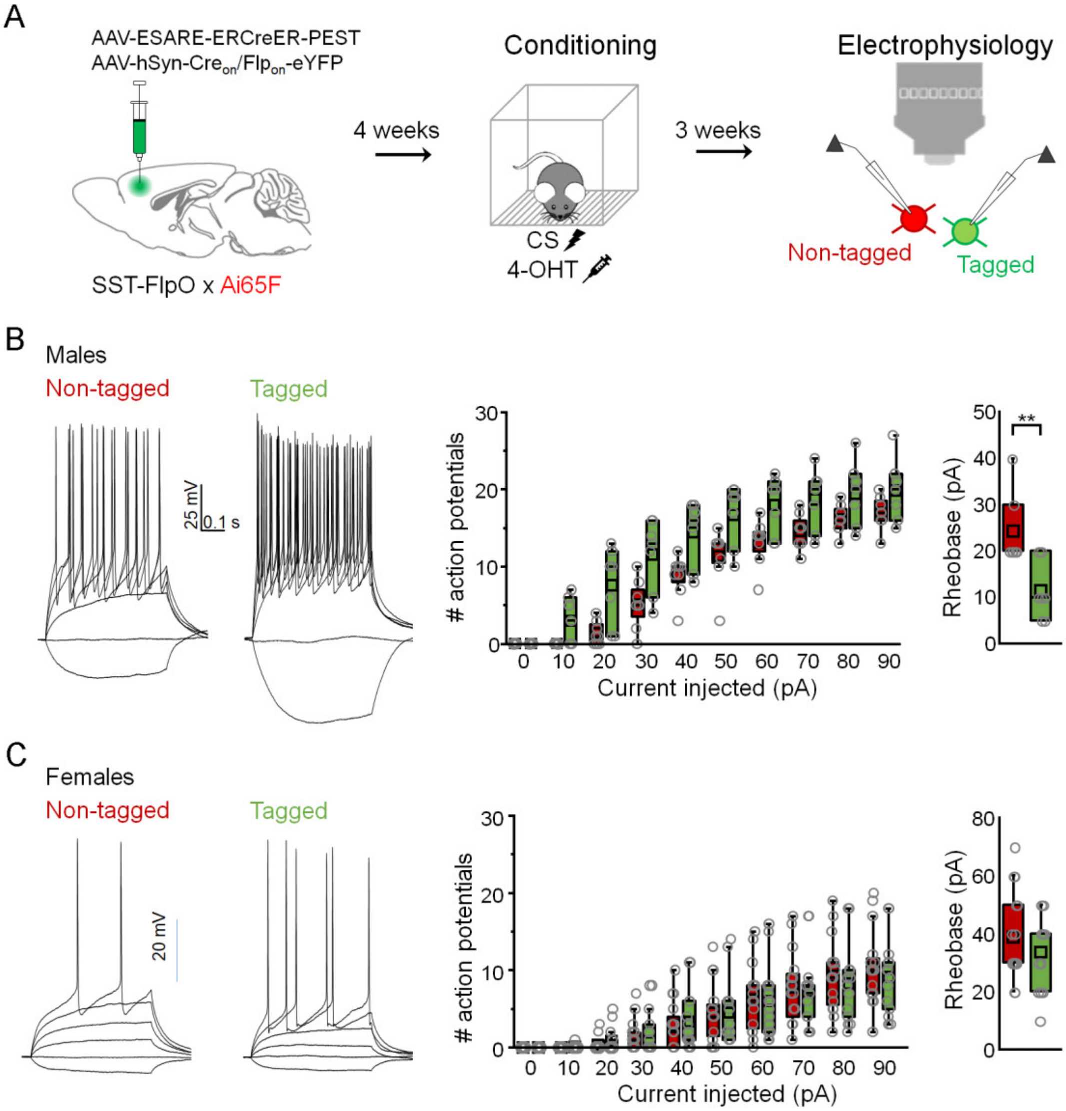


**Figure S9, related to Figure 6. Comparison of intrinsic excitability between tagged and non-tagged SST-INs of male and female mice after fear conditioning. (A)** SST-FlpO/ Ai65F double transgenic mice received prelimbic infusions of a cocktail of vectors encoding E-SARE-ERCreER, as well as Cre- and Flp-dependent eYFP. Mice were subjected to CS-US pairing and were immediately injected with 4-hydroxytamoxifen (4-OHT). Three weeks later, recordings were obtained from eYFP+/ tdTomato+ (tagged) and eYFP-/ tdTomato+ (non-tagged) SST-INs. **(B)** Representative spike trains (-20 pA and +10-40 pA current injections), input-output curves, and rheobase quantification for tagged (n = 7 cells) and non-tagged (n = 7 cells) SST-INs in the same slices of male mice (n = 3 slices from 3 mice). Input-output: $F_{(9,36)} = 1.81$, $p = 0.1$, 2-way repeated measures ANOVA. Rheobase: U = 44, $p = 0.009$, Mann-Whitney U-test. **(C)** Representative spike trains (-20 pA and +10-40 pA current injections), input-output curves, and rheobase quantification for tagged (n = 14 cells) and non-tagged (n = 16 cells) SST-INs in the same slices of female mice (n = 3 slices from 3 mice). Input-output: $F_{(9,99)} = 0.522$, $p = 0.855$, 2-way repeated measures ANOVA. Rheobase: U = 126, $p = 0.564$, Mann-Whitney U-test. **, $p < 0.01$ by Mann-Whitney U-test (**B**: rheobase).

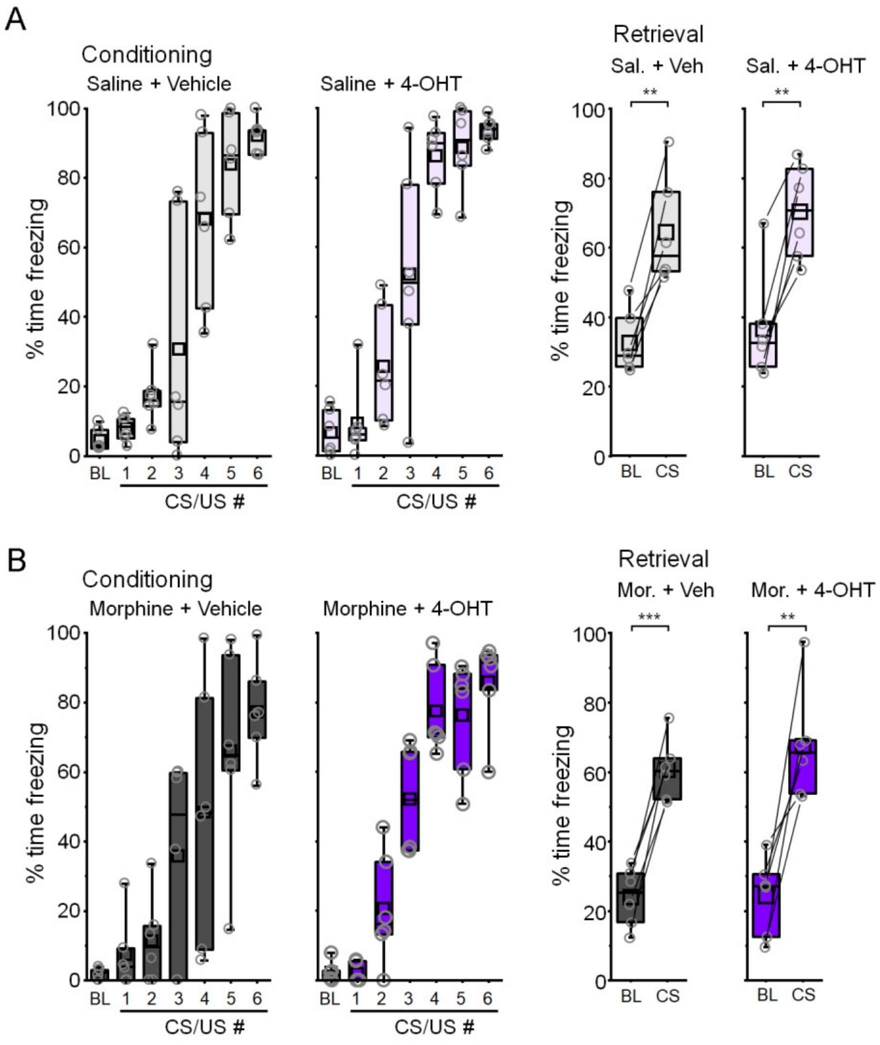


**Figure S10, related to Figure 7. Quantification of freezing for morphine-treated mice in Fig 7. (A)** Freezing during conditioning and memory retrieval for mice injected with saline. Saline vehicle retrieval: $t_5 = -5.94$, $p = 0.002$, paired t-test, n = 6 mice. Saline 4-OHT retrieval: $t_5 = -5.94$, $p = 0.002$, paired t-test, n = 6 mice. **(B)** Freezing during conditioning and memory retrieval for mice injected with morphine. Morphine vehicle retrieval: $t_5 = -9.35$, $p = 2.36 \times 10^{-4}$, paired t-test, n = 6 mice. Morphine 4-OHT retrieval: $t_5 = -5.4$, $p = 0.003$, paired t-test, n = 6 mice. Experiment was performed in 2 different cohorts and pooled together. **, $p < 0.01$, ***, $p < 0.001$ by paired t-test.

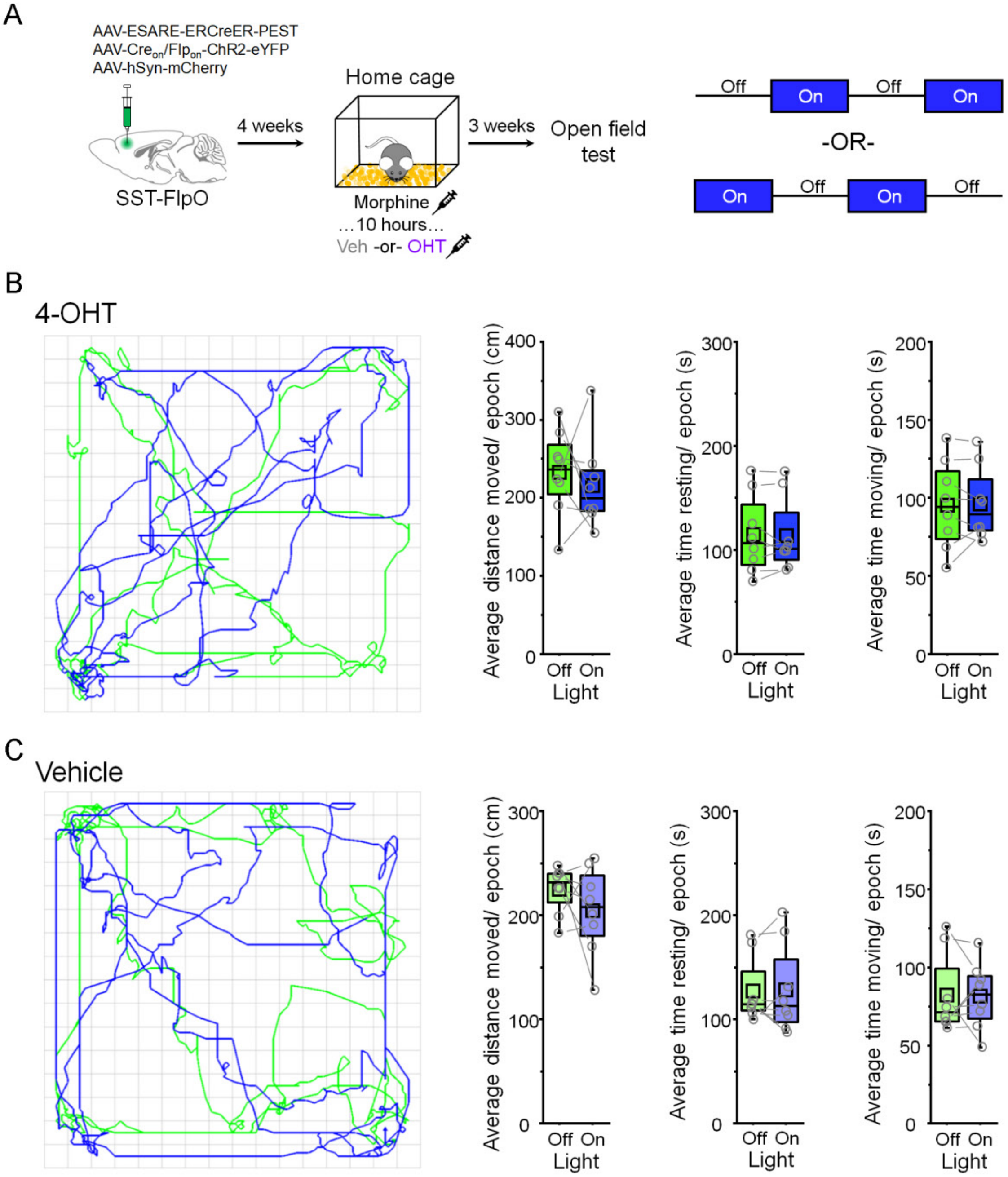


**Figure S11, related to Figure 8. Photostimulation of morphine-activated SST-INs does not affect locomotion. (A)** SST-FlpO transgenic mice received bilateral prelimbic infusions of a cocktail containing vectors encoding E-SARE-ERCreER, Cre- and Flp-dependent ChR2, and hSyn-mCherry. Four weeks later, mice were injected with morphine (10 mg/kg) followed 10 hours later by injections of vehicle (veh) or 4-hydroxytamonifen (4-OHT). After three weeks, locomotion was tested in a 20-minute open field test during light-on and light-off periods (473 nm, 10 ms pulses, 20 Hz). Light on/off epochs were 5 minutes long and were presented in a counterbalanced fashion. **(B)** Example activity plot and quantification of locomotor parameters for 4-OHT mice (n = 8 mice). Distance moved: $t_7 = 0.639$, $p = 0.543$, paired t-test. Time resting: $W = 16$, $p = 0.833$, Wilcoxon signed rank test. Time moving: $t_7 = -0.159$, $p = 0.878$, paired t-test. **(C)** Example activity plot and quantification of locomotor parameters for vehicle mice (n = 8 mice). Distance moved: $t_7 = 1.14$, $p = 0.292$, paired t-test. Time resting: $W = 18$, $p = 1$, Wilcoxon signed rank test. Time moving: $W = 18$, $p = 1$, Wilcoxon signed rank test. Experiments were performed in 2 different cohorts and pooled together.
